# Functional divergence of the NSs protein defines interferon antagonism and viral fitness across Bhanja virus lineages

**DOI:** 10.64898/2026.06.08.730832

**Authors:** Andrew T. Clarke, Kelsey Davies, Mazigh Fares, Agnieszka M. Szemiel, Diogo Correa Mendonca, Karen Kerr, Vanessa Herder, Lesley Bell-Sakyi, Brian J. Willett, Arvind H. Patel, Alain Kohl, Benjamin Brennan

## Abstract

Bhanja virus (BHAV) is a tick-borne bandavirus (Family *Phenuiviridae*) with a broad geographic distribution and documented neuroinvasive capacity, yet its molecular biology remains poorly understood. Within the genus *Bandavirus*, the non-structural protein NSs functions as the primary antagonist of innate immune responses, although mechanistic details for BHAV remain unknown. We established a reverse genetics platform for BHAV by combining virion RNA sequencing with terminal untranslated region mapping, enabling generation of recombinant viruses and systematic investigation of viral determinants. Using this system, we characterized recombinant BHAV replication in mammalian and arthropod cell lines and demonstrated that interferon competence is a critical determinant of viral replication in mammalian cells. Notably, we observed substantial amino acid divergence within the NSs protein across geographically distinct BHAV isolates, despite higher conservation of other viral proteins. To investigate the functional significance of this variation, we generated recombinant viruses expressing heterologous NSs proteins from African and European isolates. Viruses expressing NSs from ibAr2709 or R1819 isolates exhibited enhanced capacity to suppress interferon-beta induction compared to the prototype IG690 strain, correlating with increased viral protein accumulation in interferon-competent cells. Mechanistic studies revealed that BHAV NSs proteins inhibit interferon induction upstream of IRF3, with ibAr2709 and R1819 NSs showing selective inhibition of TBK1 phosphorylation. However, unlike highly pathogenic bandaviruses such as severe fever with thrombocytopenia syndrome virus (SFTSV), BHAV NSs proteins exhibit comparatively weak antagonism of downstream interferon signalling. *In vivo* studies using interferon alpha/beta receptor-deficient mice demonstrated that NSs sequence variation influences viral replication in splenic tissue without substantially altering disease phenotype. Together, these findings establish BHAV reverse genetics tools for future investigation and reveal how naturally-occurring NSs divergence modulates innate immune antagonism and viral fitness while maintaining an overall attenuated disease phenotype.

**Author Summary:** Bhanja virus is a tick-borne virus distributed across Asia, Africa, and Europe that causes neurological disease, yet little is known about how it replicates or evades immunity. We developed the laboratory tools to create recombinant Bhanja viruses and investigate viral functions. We discovered that the virus replicates efficiently in cells lacking interferon responses but is severely restricted in normal immune-competent cells, revealing interferon as a major barrier to viral growth. We observed that Bhanja virus isolates from distinct geographic regions have different versions of a viral protein called NSs that suppresses interferon responses. Isolates from Africa and Europe were better at blocking interferon production than the original Asian strain, correlating with increased viral growth in immune-competent cells. However, unlike related viruses that cause more serious disease such as SFTSV, BHAV NSs proteins only partially antagonise interferon. While this weak antagonism likely contributes to the attenuated phenotype observed in our experimental systems, the natural pathogenesis of Bhanja virus in human hosts and tick vectors remains to be determined. This study provides tools for examining Bhanja virus biology and demonstrates that small viral differences in viral sequence can meaningfully influence evasion of innate immunity and viral fitness without triggering severe disease.

## INTRODUCTION

Bhanja virus (ICTV: *Bandavirus bhanjanagarense*, BHAV) is a neurotropic tick-borne pathogen belonging to the family *Phenuiviridae* within the order *Bunyavirales*. BHAV was first isolated in India in 1954 from a pool of *Haemaphysalis intermedia* ticks attached to a goat presenting with paralysis [1]. Since then, BHAV has been isolated across central Asia, sub-Saharan Africa and central Europe from multiple tick genera, including *Haemaphysalis*, *Dermacentor*, *Hyalomma*, *Amblyomma* and *Rhipicephalus* [1–9]. Isolation of BHAV from vertebrates is rare, however high seropositivity has been detected throughout endemic regions in domestic animals including cattle, sheep, goats and dogs [10–15]. Experimental BHAV infection in young ruminants and rodents can cause fever, leukopenia and meningoencephalitis [16–20]. In contrast, adult animals generally remain asymptomatic following subcutaneous (s.c.) or intravenous (i.v.) inoculation yet succumb to disease when virus is administered intranasally (i.n.) or intracranially (i.c.) [17,18,20]. These findings indicate that BHAV exhibits a neurotropic phenotype, with pathogenesis strongly dependent on route of inoculation and the age of the host. Human infection is considered rare, with only one confirmed naturally-acquired case reported alongside several laboratory-acquired infections [21–23]. Clinical disease is typically febrile and characterised by headache, arthralgia, myalgia and conjunctivitis, although progression to meningoencephalitis with vomiting, photophobia and partial paralysis has also been documented [21,22,24]. Despite this, BHAV likely remains substantially under-recognised, particularly within endemic regions of the Balkans where serological surveys have demonstrated unexpectedly high levels of exposure [15,25,26]. Supporting this, recent studies identified serological evidence of recent BHAV infection in Croatian patients presenting with neuroinvasive disease of unknown aetiology [27].

Phylogenetic analyses place BHAV within the genus *Bandavirus*, where it forms a distinct lineage closely related to several medically important tick-borne bandaviruses, including severe fever with thrombocytopenia syndrome virus (ICTV: *Bandavirus dabieense*, SFTSV) and Heartland virus (ICTV: *Bandavirus heartlandense*, HRTV) [28]. Since its emergence, SFTSV has caused more than 16,000 reported human infections across East Asia, with case fatality rates ranging from 5.2% to 32.6% [29–31]. Clinically, SFTSV infection can present with thrombocytopenia, leukopenia, haemorrhagic manifestations, neurological complications and multi-organ failure [32]. Similarly, HRTV has emerged as a cause of severe tick-borne febrile illness in North America, with reported cases presenting with fever, fatigue, leukopenia and thrombocytopenia [33,34]. Despite belonging to the same genus as these highly pathogenic viruses, the molecular virology, pathogenicity and host-pathogen interactions of BHAV remain poorly understood.

The BHAV genome consists of three single-stranded RNA segments designated small (S), medium (M) and large (L). Collectively, these encode the nucleocapsid protein (N), RNA-dependent RNA polymerase (RdRp), glycoprotein precursor (Gn/Gc) and a non-structural protein (NSs). As with other phenuiviruses, the S segment utilises an ambisense coding strategy whereby the N protein is translated directly from the genomic S RNA, while NSs is expressed from a subgenomic mRNA transcribed from the antigenomic replication intermediate [35]. The untranslated terminal regions of each segment serve as promoters for viral transcription and replication through formation of ribonucleoprotein (RNP) complexes involving N, RdRp and the viral genomic RNA.

Within the *Bandavirus* genus, the NSs protein functions as the major virulence factor and principal antagonist of innate immune responses [36]. Despite exhibiting limited amino acid conservation, NSs proteins from different bandaviruses have evolved mechanistically distinct strategies to suppress antiviral signalling. SFTSV NSs forms prominent cytoplasmic inclusion bodies that spatially sequester multiple innate immune signalling proteins including TBK1, IRF3 and STAT family members, thereby suppressing both interferon induction and signalling pathways [37–41]. In contrast, HRTV NSs displays diffuse cytoplasmic localisation and antagonises interferon responses through a distinct TBK1-centred mechanism without the formation of large inclusion body structures [42,43]. These observations suggest substantial mechanistic diversity in the modulation of host antiviral responses by bandaviral NSs proteins. However, the mechanisms by which BHAV NSs antagonises innate immunity remain entirely unknown.

Although relatively few BHAV genome sequences are currently available, existing data indicate substantial divergence within the NSs coding region between isolates originating from Asia, Africa and Europe [28]. This degree of variation is notable given the comparatively high conservation observed across other viral proteins, including N, and raises the possibility that naturally occurring BHAV lineages may differ in virulence and in their capacity to modulate host antiviral responses. Whether such divergence alters viral fitness, innate immune antagonism or pathogenesis has not been fully investigated [28].

Reverse genetics systems have become essential tools for dissecting bunyavirus replication, virulence and host interactions [44–51]. However, despite the broad geographic distribution and potential neuroinvasive capacity of BHAV, no recombinant system currently exists for this virus. Consequently, fundamental aspects of BHAV biology remain unresolved, including the determinants of mammalian and arachnid cell tropism, the contribution of NSs to innate immune antagonism, and whether naturally occurring NSs variation influences viral replication or pathogenesis.

In this study, virion RNA sequencing and terminal untranslated region mapping were combined to establish the first reverse genetics platform for BHAV. Using these tools, authentic recombinant BHAV were generated alongside recombinant viruses expressing heterologous NSs proteins derived from geographically distinct isolates. Subsequently, BHAV replication was evidenced in both mammalian and arachnid cell systems, defining the contribution of divergent NSs proteins to interferon antagonism and viral fitness, and investigating the underlying mechanisms governing innate immune evasion. Together, these findings provide novel insights into the molecular biology of this neglected tick-borne bandavirus and establish a framework for future investigations into BHAV replication, pathogenesis and host adaptation.

## RESULTS

### Genome characterisation of Bhanja virus IG690

Bhanja virus (BHAV) was first isolated in India in 1954 from a pool of *Haemaphysalis intermedia* ticks collected from a paralysed goat, yielding the prototype strain IG690. As the parental virus used throughout this study and one of the few BHAV isolates for which genomic data are available, IG690 was subjected to deep sequencing to validate its genome sequence and define the terminal untranslated regions required for reverse genetics system development [9,24].

Next-generation sequencing of viral RNA derived from virions isolated from a wild-type virus preparation (wtIG690) generated a high-depth dataset that enabled accurate reconstruction of the viral genome. A total of 1,091,142 reads (35–151 bp) were obtained and mapped to reference IG690 genome segments, with strong coverage across all three segments. Specifically, 34,935 reads aligned to the S segment (99.5% identity), 116,246 reads to the M segment (99.6% identity), and 134,213 reads to the L segment (99.1% identity), supporting a high-confidence consensus sequence. Genome coverage analysis revealed complete breadth across all three genomic segments (100% coverage), with mean read depth of 1,755X (median 1,178X, range 1–10,620X; Table 1). Segment-specific coverage was: S segment (mean 1,327X, 100% breadth), M segment (mean 2,501X, 100% breadth), and L segment (mean 1,493X, 100% breadth). Read depth was uniformly high across coding regions, although reduced coverage (<50 reads) was observed at the 5′ and 3′ untranslated regions, indicating incomplete resolution of genome termini and the need for targeted approaches such as RACE to define these sequences (Fig 1A).

**Fig 1.**
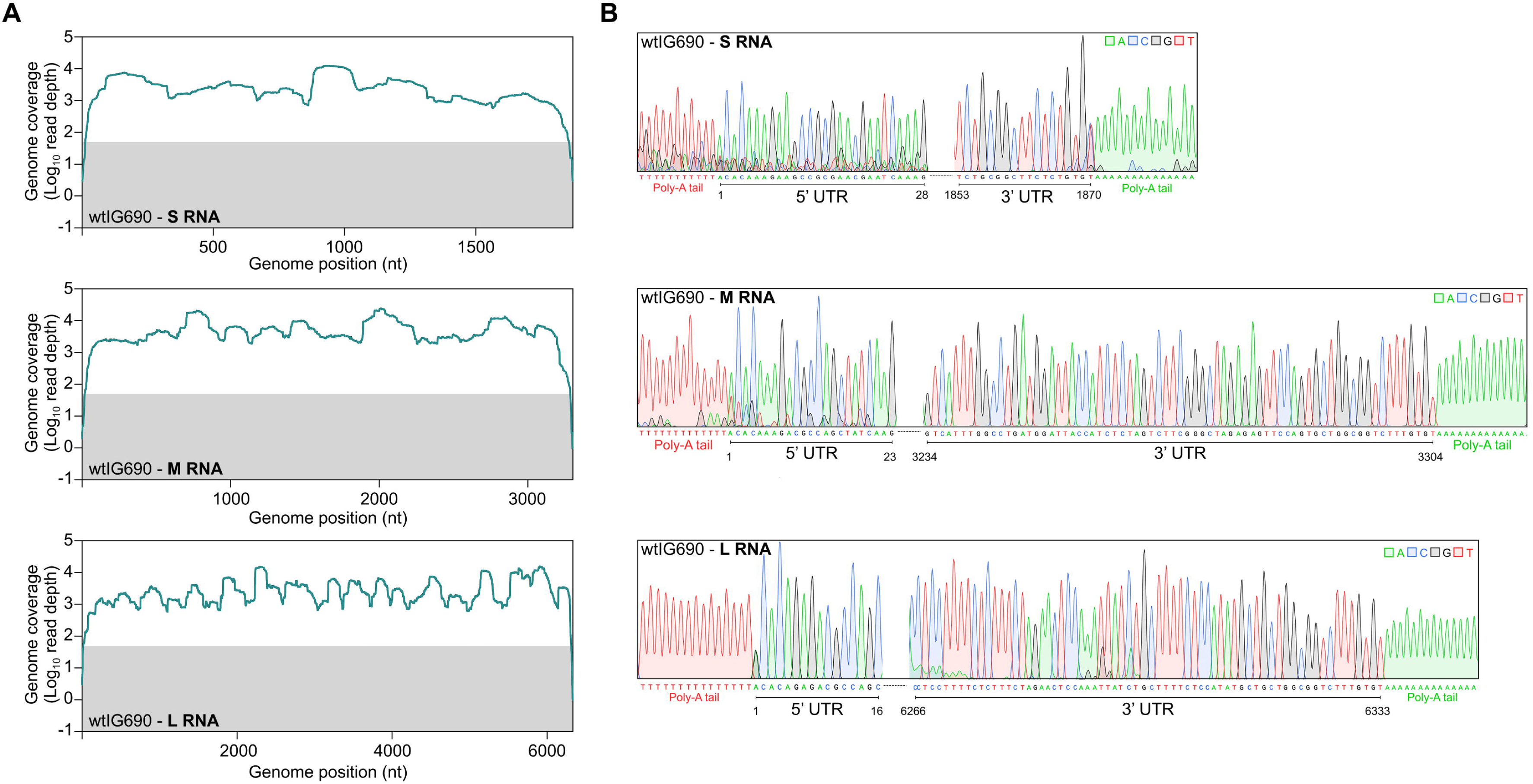
RNA sequencing and terminal untranslated region analysis of Bhanja virus wtIG690 genomic RNA segments. **(A)** RNA sequencing read coverage across the antigenomic S, M and L genome segments of wtIG690. RNA sequencing reads derived from purified wtIG690 virion RNA were aligned to the published wtIG690 antigenomic reference sequences for the S (JX961621.1), M (JX961620.1) and L (JX961619.1) segments described by Matsuno et al., 2013. Read coverage is shown in light blue and represents the Log_10_ number of sequencing reads mapped to each nucleotide position on the reference sequence. The grey shaded region indicates positions with fewer than 50 mapped reads. **(B)** Chromatographs showing the nucleotide sequences of the 5′ and 3′ untranslated regions (UTRs) of the wtIG690 S, M and L genome segments. Viral RNA purified from wtIG690 virions was subjected to 3′ rapid amplification of cDNA ends (3′RACE) prior to sequence analysis. Poly(A) tails introduced during the 3′RACE procedure are indicated.

**Table 1.**
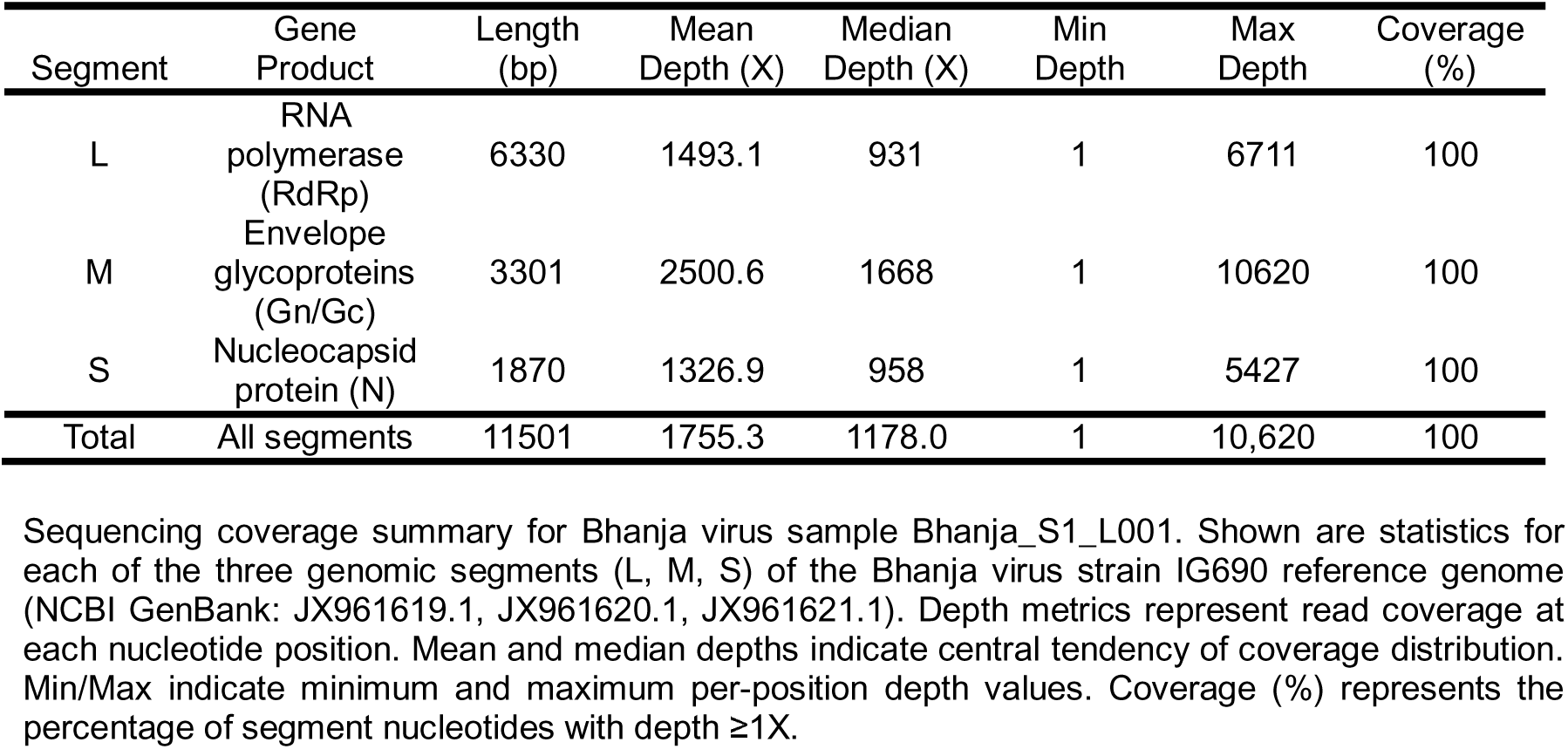
Bhanja virus genome coverage summary by sequencing segment.

The wtIG690 consensus sequences were compared with two published reference genomes [28,52] to assess sequence concordance. Across all segments, limited nucleotide differences were observed (Table 2). At 11 positions, the consensus matched the previously published IG690 reference sequence reported by Matsuno et al. [28], corresponding to four amino acid residues within the N, Gn and L proteins. Conversely, five positions aligned with the alternative IG690 sequence reported by Gmyl et al. [52], resulting in two amino acid residues within the N and Gc proteins that differed from the Matsuno reference. Overall, the consensus sequence was highly conserved, differing by only a single unique nucleotide substitution in the Gn coding region (98T>A), which resulted in a conservative amino acid change.

**Table 2.**
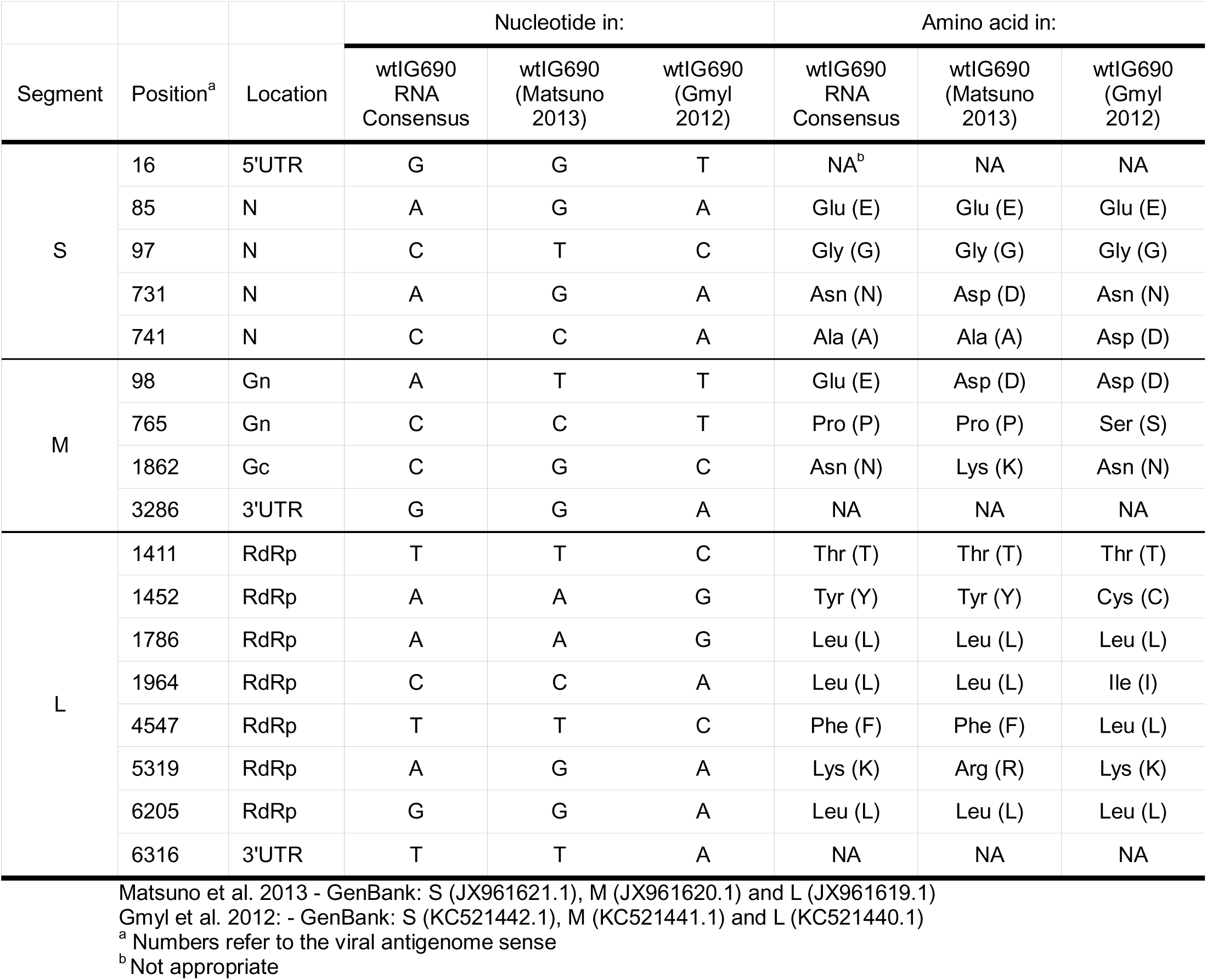
Nucleotide and amino acid changes between virus stock of Bhanja virus wtIG690 utilised in this study and two published wtIG690 sequences.

| Segment | Position<sup>a</sup> | Location | Nucleotide in: |  |  | Amino acid in: |  |  |
| --- | --- | --- | --- | --- | --- | --- | --- | --- |
|  |  |  | wtIG690<br>RNA<br>Consensus | wtIG690<br>(Matsuno<br>2013) | wtIG690<br>(Gmyl<br>2012) | wtIG690<br>RNA<br>Consensus | wtIG690<br>(Matsuno<br>2013) | wtIG690<br>(Gmyl<br>2012) |
| S | 16 | 5'UTR | G | G | T | NA<sup>b</sup> | NA | NA |
|  | 85 | N | A | G | A | Glu (E) | Glu (E) | Glu (E) |
|  | 97 | N | C | T | C | Gly (G) | Gly (G) | Gly (G) |
|  | 731 | N | A | G | A | Asn (N) | Asp (D) | Asn (N) |
|  | 741 | N | C | C | A | Ala (A) | Ala (A) | Asp (D) |
| M | 98 | Gn | A | T | T | Glu (E) | Asp (D) | Asp (D) |
|  | 765 | Gn | C | C | T | Pro (P) | Pro (P) | Ser (S) |
|  | 1862 | Gc | C | G | C | Asn (N) | Lys (K) | Asn (N) |
|  | 3286 | 3'UTR | G | G | A | NA | NA | NA |
| L | 1411 | RdRp | T | T | C | Thr (T) | Thr (T) | Thr (T) |
|  | 1452 | RdRp | A | A | G | Tyr (Y) | Tyr (Y) | Cys (C) |
|  | 1786 | RdRp | A | A | G | Leu (L) | Leu (L) | Leu (L) |
|  | 1964 | RdRp | C | C | A | Leu (L) | Leu (L) | Ile (I) |
|  | 4547 | RdRp | T | T | C | Phe (F) | Phe (F) | Leu (L) |
|  | 5319 | RdRp | A | G | A | Lys (K) | Arg (R) | Lys (K) |
|  | 6205 | RdRp | G | G | A | Leu (L) | Leu (L) | Leu (L) |
|  | 6316 | 3'UTR | T | T | A | NA | NA | NA |

Matsuno et al. 2013 - GenBank: S (JX961621.1), M (JX961620.1) and L (JX961619.1)
Gmyl et al. 2012: - GenBank: S (KC521442.1), M (KC521441.1) and L (KC521440.1)
<sup>a</sup> Numbers refer to the viral antigenome sense
<sup>b</sup> Not appropriate

Given prior reports of incorrect or incomplete UTR sequences in related bunyavirus sequences impairing minigenome activity and virus rescue [46,50,53], the wtIG690 UTRs were validated by 3′ RACE. Viral RNA was polyadenylated, reverse transcribed using an oligo(dT) primer, and amplified using segment-specific primers (SI Appendix, S1 Table). Sanger sequencing yielded high-quality chromatograms spanning the genome termini. All UTR sequences showed complete identity to the Matsuno reference and differed from the Gmyl sequence at three positions (Fig 1B).

To define transcription termination within the wtIG690 S segment, 3′ RACE was performed on RNA from infected cells. Sanger sequencing revealed that both N and NSs mRNAs terminate within the intergenic region, at nucleotide positions 888 and 850, respectively. These data clarify the location of termination signals and define the correct S segment transcriptional organisation (SI Appendix, S1 Fig.).

Overall, these data confirm the high sequence fidelity of the laboratory wtIG690 stock relative to published reference genomes, supporting its use as a robust template for reverse genetics system development.

### Recombinant IG690 is rescued efficiently and recapitulates wild-type replication and interferon sensitivity

Consistent with the high sequence fidelity of the wtIG690 stock, a recombinant IG690 virus (rIG690) was rescued successfully using a plasmid-based reverse genetics system, as described previously for related bunyaviruses (Fig. 2) [46,50]. Infectious virus was recovered following co-transfection of BSR-T7/5 cells, with rIG690 generated reproducibly at a mean titre of 9.76 × 10^2^ focus-forming units (FFU)/ml across two independent rescue experiments.

**Fig 2.**
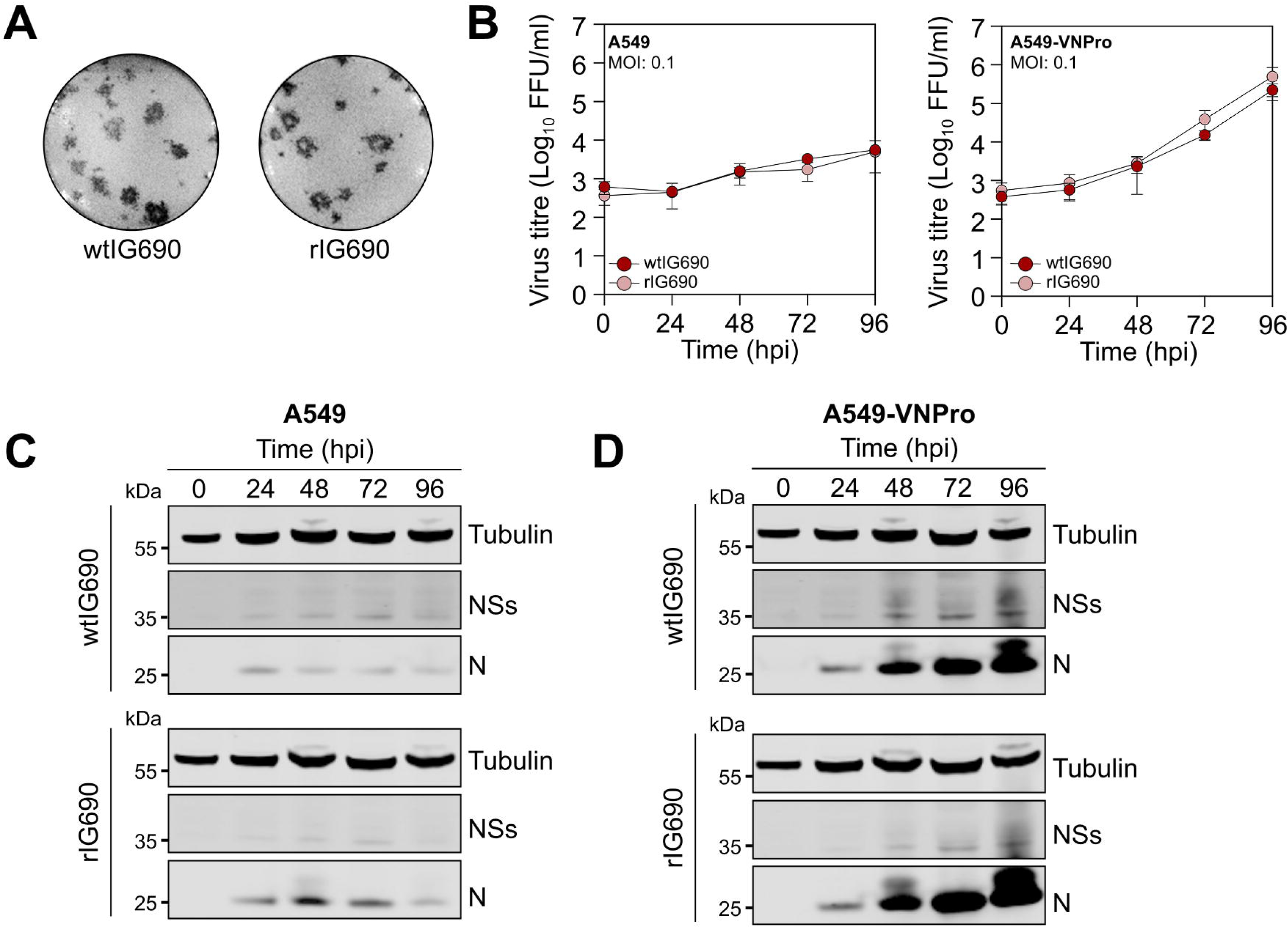
Recombinant Bhanja virus IG690 is efficiently rescued and recapitulates wild-type replication and interferon sensitivity. **(A)** Representative immunofocus assay images showing viral foci produced by parental wtIG690 and recombinant rIG690 following infection of Vero E6 cell monolayers. **(B)** Replication kinetics of wtIG690 and rIG690 in A549 and A549-VNPro cells. Cell monolayers were infected at an MOI of 0.1 FFU/cell and culture supernatants harvested at the indicated times p.i. for titration by focus-forming assay on Vero E6 cells. Viral titres are presented as Log_10_ FFU/ml. Data represent the mean ± SD from three biological replicates. **(C,D)** Representative analysis of viral protein expression during wtIG690 and rIG690 infection of A549 **(C)** and A549-VNPro **(D)** cells. Cell lysates collected at the indicated times p.i. from growth curves described in **(B)** were analysed by western blotting for expression of BHAV N and NSs proteins. Tubulin was used as a loading control. Representative blots from the three independent experiments are shown.

Foci formed in immunofocus assays were comparable in morphology to those of the parental wtIG690 isolate, and viral stocks were subsequently amplified to generate working stocks for downstream experimentation (Fig. 2A).

Comparative analysis in type I interferon-competent (A549) and IFN-deficient (A549-VNPro) cells showed that rIG690 closely recapitulates the growth kinetics of wtIG690 (Fig. 2B). In A549 cells, both viruses exhibited restricted replication, with titres increasing modestly from ∼10^2^–10^3^ FFU/ml at early time points to ∼10^4^ FFU/ml by 96 h post-infection (p.i.). In contrast, in A549-VNPro cells, replication was markedly enhanced, with both wtIG690 and rIG690 reaching titres approaching ∼10^6^ FFU/ml by 96 h p.i., demonstrating equivalent sensitivity to type-I interferon restriction.

Consistent with these growth data, viral protein expression profiles were comparable between wtIG690 and rIG690 (Fig. 2C,D). In both cases, N and NSs proteins accumulated progressively over the course of infection, with minimal N expression at early time points and robust detection from 48–96 h p.i., with stronger N signal observed in BHAV-infected A549-VNPro cells. NSs expression was comparable between wtIG690 and rIG690 in both cell types, although levels were higher and detected earlier in A549-VNPro cells than in A549 cells. Together, these data confirm that rIG690 mirrors the parental isolate in both replication kinetics and temporal protein expression.

### Cell-type permissibility reveals interferon restriction as a key determinant of BHAV replication

To identify cell systems that support efficient BHAV replication, a panel of mammalian cell lines from diverse host species and tissues was screened for permissibility to wtIG690 infection (Fig 3). The cell lines were infected at MOI 0.01 and 0.1 FFU/cell, and viral titres and N protein expression were assessed at 5 days post-infection. Replication efficiency varied markedly across cell types (Fig. 3A), with the highest titres observed in cells deficient in type-I interferon responses, including Vero E6, BHK-21, Huh7 and Huh7.5 cells, where titres typically reached ∼10^6^–10^7^ FFU/ml. In contrast, replication was restricted strongly in immune-competent A549 cells, with titres remaining at ∼10^2^-10^3^ FFU/ml and coupled with minimal N protein expression (Fig. 3B). In A549-derived cells with disrupted innate immune signalling (A549-NPro and A549-VNPro), virus replication was restored, as evidenced by virus titres achieving ∼10^6^–10^7^ FFU/ml and robust N protein accumulation. In these cell lines, innate immune responses are suppressed through constitutive expression of the bovine viral diarrhoea virus (BVDV) NPro protein, which targets IRF3 for proteasomal degradation, either alone (A549-NPro; [54]) or in combination with the parainfluenza virus 5 (PIV5) V protein, which antagonises STAT1-dependent interferon signalling (A549-VNPro; [55]).

**Fig 3.**
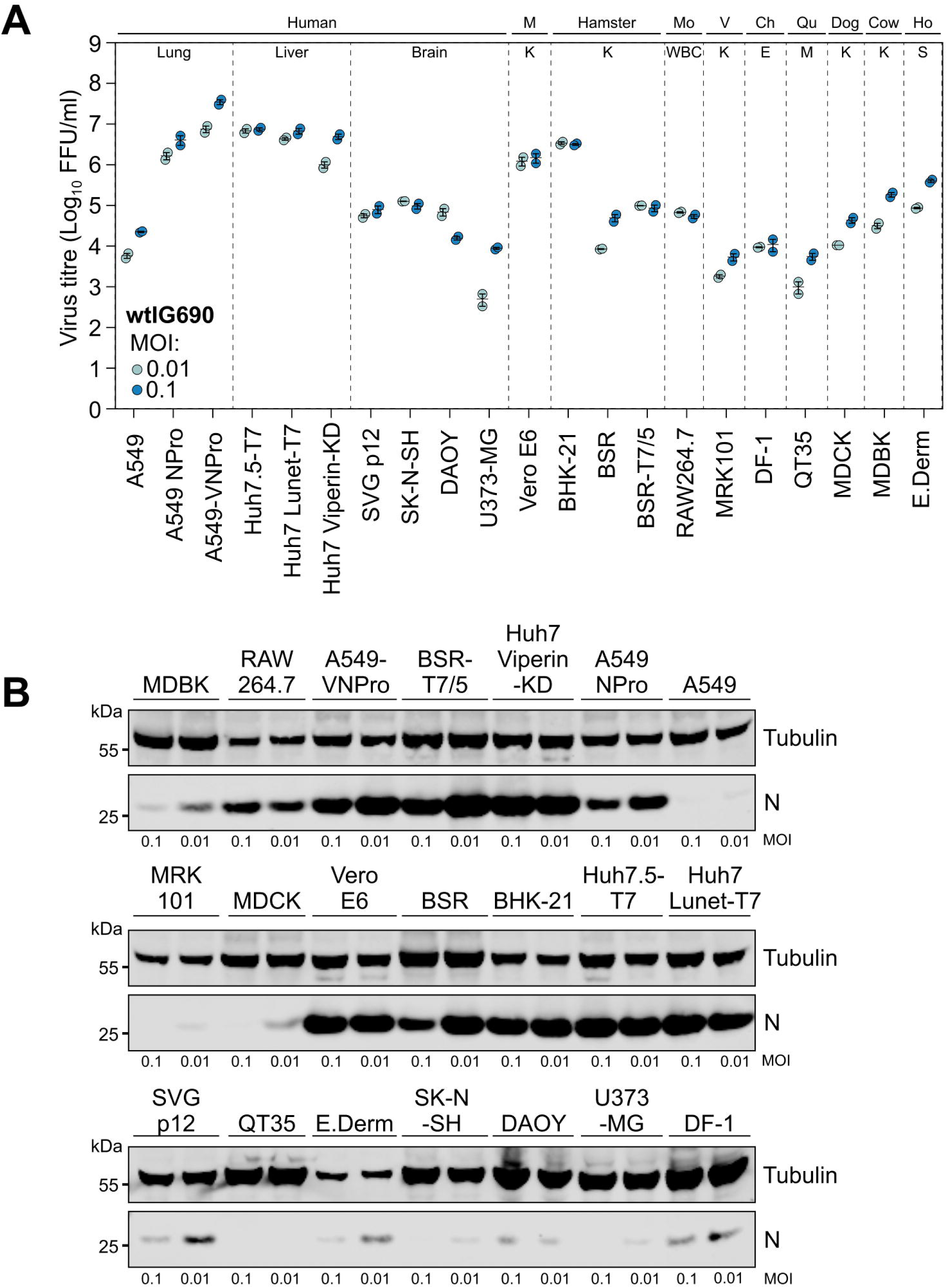
Cell-type permissibility reveals interferon restriction as a key determinant of Bhanja virus (BHAV) replication. **(A)** Replication of wtIG690 in a panel of vertebrate cell lines derived from different host species and tissues. Cell monolayers were infected with wtIG690 at an MOI of 0.01 FFU/cell (light blue) or 0.1 FFU/cell (dark blue). At 5 d p.i., culture supernatants were harvested and viral titres determined by focus-forming assay on Vero E6 cells. Viral titres are presented as Log_10_ FFU/ml. M, monkey; Mo, mouse; V, vole; Ch, chicken; Qu, quail, Ho, horse; WBC, white blood cells; K, kidney; E, embryo; S, skin. Experiments were performed in duplicate (n = 2). **(B)** Representative analysis of BHAV N protein expression in infected cell monolayers from the experiment shown in **(A)**. Cell lysates harvested at 5 d p.i. were analysed by western blotting using BHAV anti-N and anti-tubulin antibodies.

Despite the neurotropic disease associated with BHAV infection, the available human neuronal cell lines exhibited low permissibility/virus replication, with titres generally <10^4^ FFU/ml. Similarly, cell lines derived from rodent (with the exception of the hamster line BHK-21) and livestock species supported limited replication, typically in the range of ∼10^3^–10^5^ FFU/ml.

These findings indicate that interferon competence is a major determinant of BHAV replication in mammalian cells, although susceptibility may vary between neuronal cell types and viral isolates. Given that BHAV is maintained in arachnid vectors that lack canonical interferon pathways, we next investigated viral replication in arthropod cell lines.

### BHAV replication is restricted in *Ixodes*-derived and *Aedes-derived* cells but maintained in *Rhipicephalus*-derived cells

We next assessed the ability of rIG690 to replicate in tick- and mosquito-derived cell lines. Replication kinetics were examined over an 18-day time course in *Ixodes* spp. cell lines (IRE/CTVM19, IRE/CTVM20 and ISE6) and compared with *Rhipicephalus microplus* cell lines (BME/CTVM6 and BME/CTVM23), representing a tick genus from which BHAV has been isolated previously [56] (SI Appendix, S2 Fig.). In *Ixodes* spp. cell lines, viral titres declined progressively from ∼10^5^–10^6^ FFU/ml at early time points to ∼10^3^–10^4^ FFU/ml by day 18 p.i., accompanied by a marked reduction in N protein expression (SI Appendix, S2A,B Fig.). In contrast, replication was sustained in *Rhipicephalus*-derived cells, with titres remaining stable at ∼10^5^–10^6^ FFU/ml throughout the time course and concomitant with consistent N protein accumulation in infected cell cultures. This difference was further supported by immunofluorescence, which showed substantially higher N staining in BHAV-infected *R. microplus* cells compared to *Ixodes* spp. cells (SI Appendix, S2C Fig).

Consistent with the published host vector species for BHAV, replication was restricted markedly in mosquito-derived cell lines. In both Aag2-AF5 (*Aedes aegypti*) and C6/36 (*Aedes albopictus*) cells, viral titres declined from ∼10^5^ FFU/ml at early time points to near or below the limit of detection by day 18 (SI Appendix, S2D Fig.), with no N protein detectable by western blotting or immunofluorescence (SI Appendix, S2E, F Fig.). The residual signal observed by immunofluorescence was also present in mock-infected controls, indicating autofluorescence rather than productive infection.

Together, these findings indicate that BHAV replication is highly context-dependent, suggesting that viral determinants may modulate host-specific fitness. As NSs is the primary interferon antagonist in mammalian-infecting bunyaviruses [57], we next investigated whether sequence divergence in NSs across BHAV isolates contributes to differential interferon antagonism.

### NSs sequence divergence modulates viral protein accumulation under interferon-competent conditions

To investigate whether viral determinants contribute to the observed differences in host-dependent replication, we examined sequence variation within the NSs protein across all currently available BHAV isolates. Pairwise comparisons of NSs amino acid sequences revealed substantial divergence between lineages, with identities ranging from ∼84% to 99% (Fig. 4A & SI Appendix, S3 Fig.), indicating marked variability within this viral protein. To assess the functional consequences of this divergence, recombinant IG690 viruses expressing heterologous NSs proteins were generated alongside an NSs-deleted control virus expressing mGreenLantern (mGL) in place of NSs, utilising the reverse genetics system described previously. As the anti-BHAV NSs antibody was generated against IG690 NSs and showed limited cross-reactivity with heterologous NSs proteins, all recombinant NSs proteins were engineered with a C-terminal V5 epitope tag to facilitate direct comparisons of expression and function between isolates. (Fig. 4B). At this stage, the closely related European isolate R1336 was excluded from further analysis due to its NSs sequence being nearly identical to R1819 (99.7% identity).

**Fig 4.**
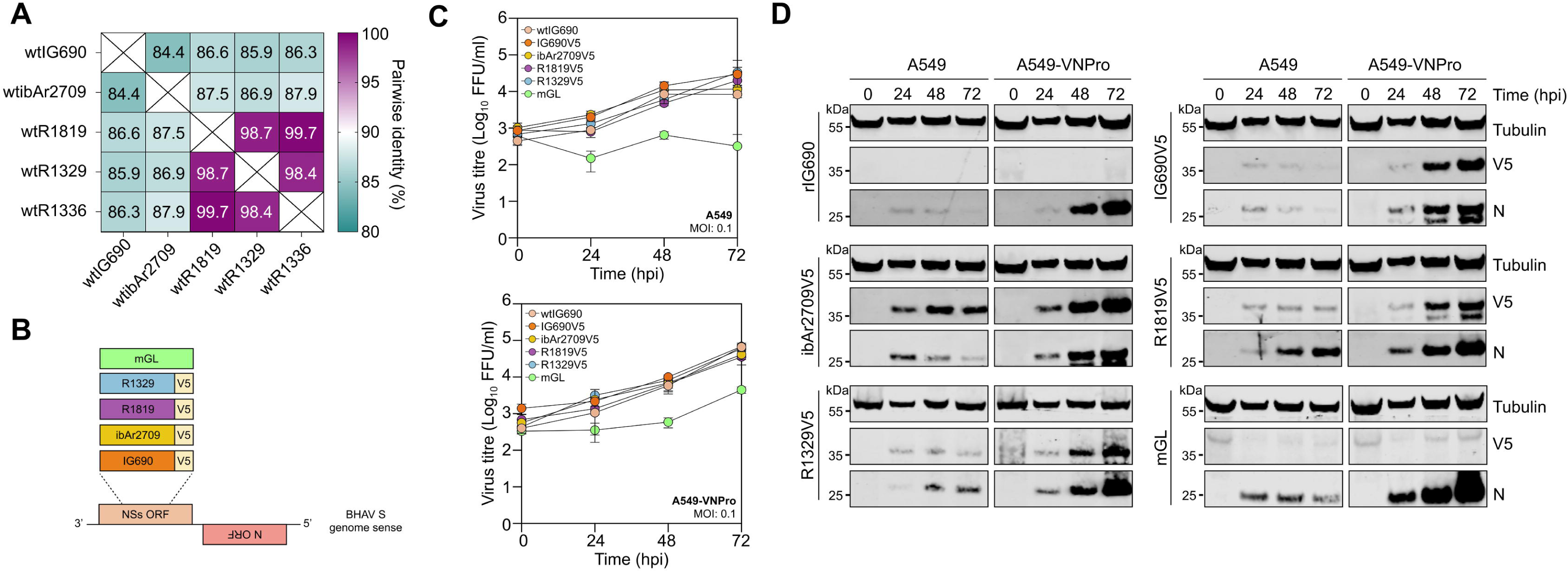
Bhanja virus (BHAV) NSs sequence divergence modulates viral protein accumulation under interferon-competent conditions. **(A)** Heat map showing pairwise amino acid identity (%) between the NSs proteins of five BHAV isolates. The following accession numbers corresponding to BHAV NSs proteins were used: IG690 (AGK25093); ibAr2709 (YP_009141016); R1819 (AGC60109); R1329 (PZ464570); R1336 (PZ464571). **(B)** Schematic representation of recombinant rIG690 viruses expressing heterologous c-terminal V5-tagged NSs proteins derived from differing BHAV isolates, generating rIG690NSsV5 (orange), ribAr2709NSsV5 (yellow), rR1819NSsV5 (purple) and rR1329NSsV5 (blue). The NSs-deletant control virus expressing mGreenLantern (mGL; green) is also shown. **(C)** Replication kinetics of recombinant BHAV expressing heterologous V5-tagged NSs proteins in A549 (top) and A549-VNPro (bottom) cells. Cell monolayers were infected at an MOI of 0.1 FFU/cell and culture supernatants harvested at the indicated times p.i. for titration by focus-forming assay on Vero E6 cells. Viral titres are presented as Log_10_ FFU/ml. Data represent the mean ± SD from three biological replicates. **(D)** Representative analysis of viral protein expression during infection with recombinant BHAV expressing heterologous V5-tagged NSs proteins. Cell lysates collected from infected A549 and A549-VNPro monolayers (described in **(C)**) at the indicated times p.i. were analysed by western blotting using anti-V5, BHAV anti-N and anti-tubulin antibodies.

Replication kinetics of these viruses were compared in IFN-competent (A549) and IFN-deficient (A549-VNPro) cells (Fig. 4C). In A549 cells, all viruses reached broadly similar titres over the course of infection, with only modest differences observed between isolates. However, viruses expressing NSs proteins from the African and European isolates ibAr2709 and R1819 exhibited increased N and NSs protein accumulation relative to wtIG690 and rIG690NSsV5 (Fig. 4D). The reduced accumulation of IG690NSsV5 relative to the heterologous NSs proteins was associated with enhanced proteasomal turnover, as treatment with the proteasome inhibitor MG132 restored IG690NSsV5 levels to those observed for ibAr2709NSsV5 and R1819NSsV5 (SI Appendix, S4 Fig.). In contrast, in A549-VNPro cells, both viral titres and viral protein expression levels were broadly comparable across all viruses, consistent with reduced type I IFN-mediated restriction.

Analysis of viral protein expression in BHAV-infected cells supported these observations, with robust expression of V5-tagged heterologous NSs proteins detected in infected cells (Fig. 4D). As expected, no V5 signal was detected for the mGL virus, confirming the absence of NSs expression in this control. Together, these data indicate that sequence divergence in BHAV NSs proteins influences viral protein accumulation under interferon-competent conditions, with limited effects on peak viral titres.

### NSs divergence modulates interferon induction without altering sensitivity to exogenous interferon

To determine whether differences in NSs sequence influence interferon antagonism, the sensitivity of wtIG690 to exogenous IFNβ was assessed. Pre-treatment of A549-NPro cells with IFNβ resulted in a strong, dose-dependent inhibition of viral replication when applied 24 h prior to infection, reducing titres by approximately 3–4 logs to ∼10^2^ FFU/ml at 1000 U (Fig. 5A). Although A549-NPro cells constitutively express the bovine viral diarrhoea virus NPro protein, which suppresses IFN induction through IRF3 degradation, downstream IFN signalling remains intact in these cells, permitting establishment of an antiviral state following exogenous IFNβ treatment [54].

**Fig 5.**
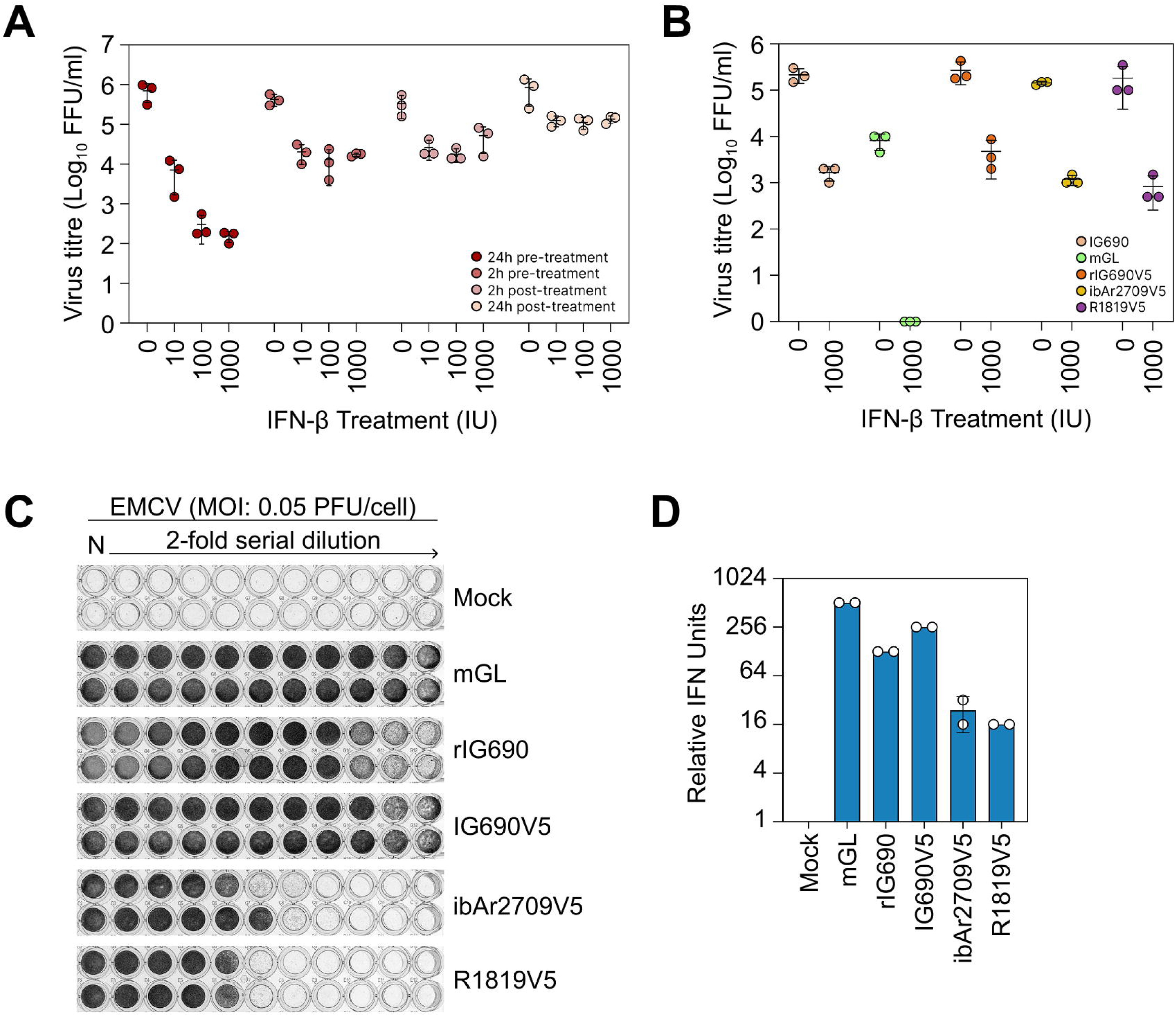
Differential interferon antagonism and IFNβ sensitivity of recombinant Bhanja virus (BHAV) expressing heterologous V5-tagged NSs proteins. **(A)** Sensitivity of rIG690 to exogenous IFNβ treatment. A549-NPro cells were treated with increasing concentrations of IFNβ (0, 10, 100 or 1000 U) either 24 h or 2 h prior to infection, or 2 h or 24 h p.i. Cells were infected with rIG690 at an MOI of 0.5 FFU/cell and culture supernatants harvested at 48 h p.i. for titration by focus-forming assay on Vero E6 cells. Viral titres are presented as Log_10_ FFU/ml. **(B)** Sensitivity of recombinant BHAV expressing heterologous V5-tagged NSs proteins to IFNβ treatment. A549-NPro cells were pre-treated with either 0 or 1000 U of IFNβ for 24 h prior to infection with recombinant viruses expressing different NSs proteins from the rIG690 backbone at an MOI of 0.5 FFU/cell. Recombinant viruses included rIG690NSsV5 (orange), ribAr2709NSsV5 (yellow), rR1819NSsV5 (purple) and rR1329NSsV5 (green). The NSs-deletant virus rIG690ΔNSs-mGL (mGL; light green) was included as a control lacking NSs expression, while rIG690 was included to assess the effect of V5-tagging on NSs function (light orange). Culture supernatants were harvested at 48 h p.i. and viral titres determined by focus-forming assay on Vero E6 cells. Viral titres are presented as Log_10_ FFU/ml. **(C,D)** Biological interferon protection assay used to assess suppression of IFN production by recombinant BHAV expressing heterologous NSs proteins. A549 cells were infected at an MOI of 2 FFU/cell with the indicated recombinant viruses. At 24 h p.i., culture supernatants were harvested, UV-inactivated and serially diluted two-fold prior to treatment of A549-NPro cells. Treated monolayers were subsequently infected with EMCV at an MOI of 0.05 PFU/cell. **(C)** Representative image of assay readout Relative IFN units are expressed as 2N, where N corresponds to the final two-fold dilution that retained protection from EMCV-induced cytopathic effect. Data shown in **(A)** and **(B)** represent the mean ± SD from three biological replicates, while data shown in **(C)** and **(D)** represent the mean ± SD from two biological replicates.

Next, the effect of NSs sequence variation on sensitivity to exogenous IFNβ was examined. In the absence of IFNβ, all viruses expressing an NSs protein replicated to comparable titres of ∼2 x10^5^ FFU/ml, whereas an NSs-deletant virus mGL was modestly attenuated (∼10^4^ FFU/ml) (Fig. 5B). Following pre-treatment with 1000 U IFNβ, titres of all NSs-expressing viruses were uniformly reduced by ∼2 logs to ∼10^3^ FFU/ml, with no clear differences between isolates. In contrast, replication of the mGL virus was ablated completely, with no detectable viral foci. Consistent with these findings, viral foci were markedly reduced in size under IFNβ treatment for all NSs-expressing viruses (SI Appendix, S5 Fig).

To directly assess the induction of IFN during infection, a biological IFN assay was performed using supernatants from infected A549 cells. As expected, mock-infected cultures produced no detectable IFN, whereas infection with the NSs-deletant mGL virus induced high levels of IFN (∼512 relative IFN units) (Fig. 5C,D). In comparison, wtIG690 (128 units) and rIG690NSsV5 (256 units) induced lower levels of IFN, while viruses expressing NSs from African and European isolates (e.g. ibAr2709, R1819 and R1329) induced IFN levels (∼16–24 units) that were reduced substantially.

Together, these data demonstrate that while NSs is required to suppress IFN induction, sequence variation between BHAV NSs proteins modulates the efficiency of this antagonism. However, this variation does not translate into differential sensitivity to exogenous IFNβ, indicating that NSs divergence primarily affects IFN induction or signalling rather than downstream antiviral restriction.

### Divergent BHAV NSs proteins inhibit interferon induction differentially at the level of TBK1/IKK**ε**

To define the step within the interferon induction pathway targeted by BHAV NSs, a plasmid-based reporter system in which individual components of the pathway were ectopically expressed was used. HEK-293T cells were co-transfected with IFNβ reporter constructs, plasmids encoding defined interferon effector proteins (RIG-I, MDA5, MAVS, TBK1, IKKε or IRF3), and NSs expression plasmids from representative BHAV isolates. SFTSV NSs, a well-established potent inhibitor of IFN induction acting at the level of TBK1 [58,59], was included as a positive control (Fig 6).

**Fig 6.**
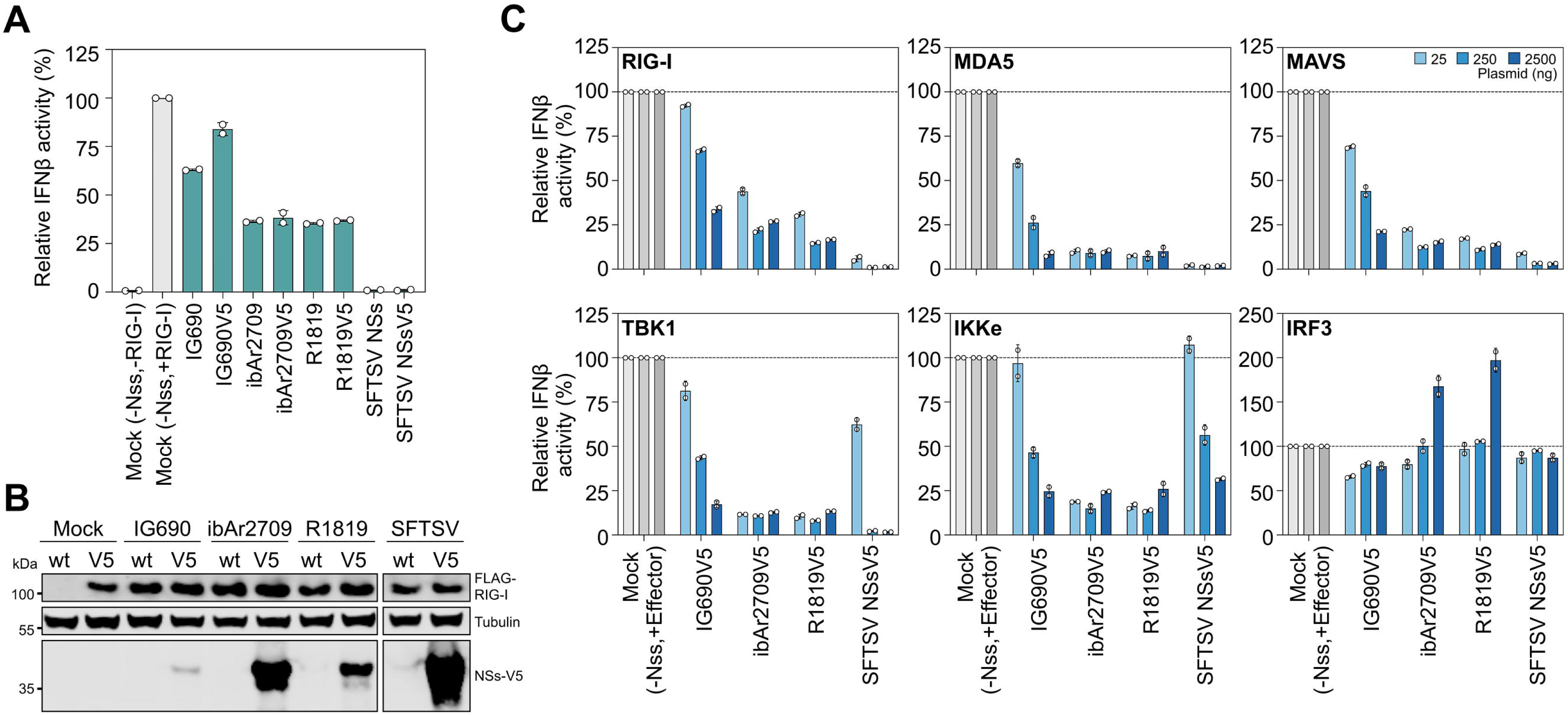
Bhanja virus (BHAV) NSs proteins differentially inhibit IFNβ induction at the level of TBK1/IKKε. **(A)** HEK-293T cells were co-transfected with p125Ffluc, pCMV-hRenilla, pCMV empty vector and 50 ng of a plasmid expressing FLAG-tagged RIG-I. Plasmids expressing either non-tagged or V5-tagged NSs proteins from BHAV isolates IG690, ibAr2709 or R1819 were co-transfected at 250 ng. V5-tagged and non-tagged SFTSV NSs constructs were included as positive controls. At 24 h post-transfection, cell lysates were harvested and firefly luciferase activity quantified. Luciferase values were normalised to hRenilla and expressed as a percentage relative to the mock control lacking NSs expression, which was set to 100% induction of the IFNβ promoter. **(B)** Representative western blot analysis of lysates from the experiment shown in **(A)**. Protein lysates were separated by SDS-PAGE and probed using anti-tubulin, anti-V5 and anti-FLAG antibodies. **(C)** HEK-293T cells were co-transfected with p125Ffluc, pCMV-hRenilla, pCMV empty vector and 50 ng of plasmids expressing the indicated effector molecules of the IFNβ induction pathway. Plasmids expressing V5-tagged BHAV NSs proteins were co-transfected at 25 ng (dark blue), 250 ng (light blue) or 2500 ng (grey). V5-tagged SFTSV NSs was included as a positive control. After 24 h incubation, cell lysates were harvested and firefly luciferase activity quantified. Luciferase values were normalised to hRenilla and expressed as a percentage relative to the mock control lacking NSs expression, which was set to 100% induction of the IFNβ promoter (dashed line). Data shown in **(A)** and **(C)** represent the mean ± SD from two independent experiments performed in duplicate.

When stimulated via RIG-I, NSs proteins from ibAr2709 and R1819 suppressed IFNβ reporter activity more efficiently than IG690 (∼25% activity), although inhibition remained less pronounced than that observed for SFTSV NSs (<10% activity) (Fig. 6A). Western blot analysis demonstrated that IG690NSsV5 accumulated at consistently lower levels than the heterologous NSs proteins under these conditions (Fig. 6B), indicating that differences in protein abundance may contribute, at least in part, to the observed variation in antagonistic activity. Reduced accumulation of IG690NSsV5 was confirmed by immunofluorescence analysis (SI Appendix, S6 Fig.). Increased plasmid input did not fully restore the inhibitory capacity of IG690NSsV5, suggesting that reduced expression alone does not account for the observed differences in IFN antagonism (SI Appendix, S4 Fig.).

Pathway dissection revealed that BHAV NSs proteins inhibited IFNβ promoter activation induced by RIG-I, MDA5 and MAVS, with ibAr2709 and R1819 showing the strongest suppression across multiple input levels (Fig. 6C). Inhibition was also observed when signalling was driven by TBK1 and IKKε, whereas IRF3-mediated activation was not suppressed, indicating that BHAV NSs acts upstream of IRF3. This pattern closely mirrors that of SFTSV NSs [37], supporting a conserved targeting of the TBK1/IKKε axis, although BHAV NSs proteins displayed reduced overall potency.

We also determined through BHAV NSs subcellular localisation analysis that similar patterns of cytoplasmic staining were observed across isolates, indicating that differences in antagonistic activity are not explained by gross differences in localisation (SI Appendix, S6 Fig.). Together, these data indicate that BHAV NSs proteins inhibit interferon induction upstream of IRF3, likely at the level of TBK1 and/or IKKε, with sequence divergence modulating the efficiency of this antagonism.

### BHAV NSs proteins do not inhibit IRF3 phosphorylation but differentially modulate TBK1 activation

To further define the mechanism by which BHAV NSs proteins inhibit interferon induction their effects on the phosphorylation of key signalling intermediates TBK1 and IRF3 were examined using a plasmid-based overexpression system. HEK-293T cells were co-transfected with RIG-I or FLAG-tagged TBK1 together with NSsV5 expression constructs, and phosphorylation of IRF3 (Ser386) and TBK1 (Ser172) was assessed by western blotting (Fig. 7A). As expected, RIG-I stimulation induced robust IRF3 phosphorylation in control conditions. In contrast, expression of BHAV NSs proteins from IG690, ibAr2709 and R1819 did not reduce IRF3 phosphorylation, although modest reductions in total IRF3 levels were observed for ibAr2709 and R1819. Increasing the expression of IG690NSsV5 did not alter this phenotype.

**Fig 7.**
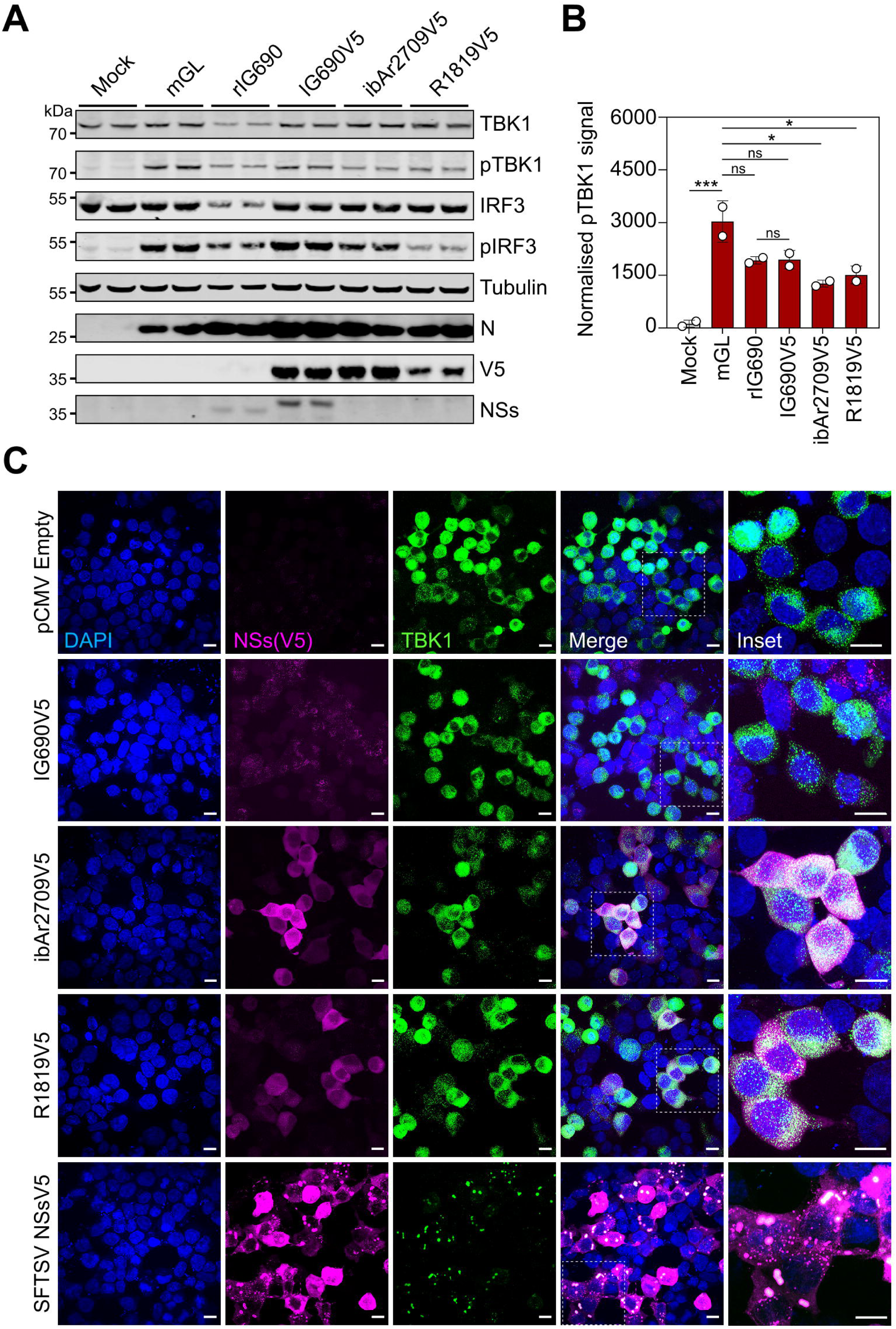
Bhanja virus (BHAV) ibAr2709 and R1819 NSs proteins selectively inhibit TBK1 phosphorylation and colocalise with TBK1. **(A)** HEK-293T cells were co-transfected with plasmids expressing FLAG-TBK1 and the indicated V5-tagged NSs proteins. SFTSV NSsV5 was included as a positive control. At 24 h post-transfection, cell lysates were harvested and analysed by western blotting using antibodies against TBK1, phosphorylated TBK1 (pTBK1), IRF3, phosphorylated IRF3 (pIRF3), BHAV N, V5, BHAV NSs and tubulin. **(B)** Quantification of phosphorylated TBK1 levels from the experiment shown in (A). Signal intensities for pTBK1 were internally normalised to tubulin. Data represent the mean ± SD from two biological replicates. Statistical significance was determined using one-way ANOVA with multiple comparisons. ns, not significant; *P < 0.05; ***P < 0.001. **(C)** Colocalization of V5-tagged BHAV NSs proteins with FLAG-TBK1 in transfected HEK-293T cells. Cells were co-transfected with plasmids expressing FLAG-TBK1 and the indicated NSsV5 proteins. SFTSV NSsV5 was included as a positive control for TBK1 sequestration into NSs-associated inclusion bodies. At 24 h post-transfection, cells were fixed and stained using anti-V5 and anti-FLAG antibodies together with DAPI. Images were acquired using confocal microscopy. Insets show enlarged regions from the merged images. Scale bars, 10 μm.

Next, the effect of BHAV NSs proteins on upstream kinase activation was assessed (Fig. 7A, B). Among the BHAV NSs proteins, only ibAr2709NSsV5 and R1819NSsV5 reduced TBK1 phosphorylation significantly, whereas IG690NSsV5 had no detectable effect. These differences were not explained by protein expression levels, as robust expression of R1819NSsV5 was observed. Consistent with these findings, immunofluorescence analysis revealed that all BHAV NSs proteins localised to cytoplasmic puncta and showed partial co-localisation with TBK1, with no major differences in subcellular distribution between isolates (Fig. 7C).

Together, these data indicate that BHAV NSs proteins do not inhibit IRF3 activation directly, but instead act upstream in the interferon induction pathway, with ibAr2709 and R1819 NSs displaying an enhanced capacity to limit TBK1 activation.

### BHAV NSs proteins inhibit interferon signalling weakly compared to SFTSV NSs

We next assessed whether BHAV NSs proteins also antagonise downstream interferon signalling by examining activation of the interferon-stimulated response element (ISRE) promoter. HEK-293T cells were co-transfected with an ISRE luciferase reporter, Renilla control plasmid, and increasing amounts of NSsV5 expression constructs, followed by stimulation with IFNβ. SFTSV NSs, a well-characterised potent inhibitor of type I interferon signalling, was included as a positive control.

In contrast to its strong suppression of IFN induction, IG690NSsV5 exerted only a modest inhibitory effect on ISRE activation, reducing reporter activity by ∼10% at the highest plasmid input (Fig. 8). R1819NSsV5 showed a more pronounced reduction of ∼40%, whereas ibAr2709NSsV5 had little or no detectable effect across the conditions tested. By comparison, SFTSV NSs robustly inhibited ISRE activation, confirming effective assay performance.

**Fig 8.**
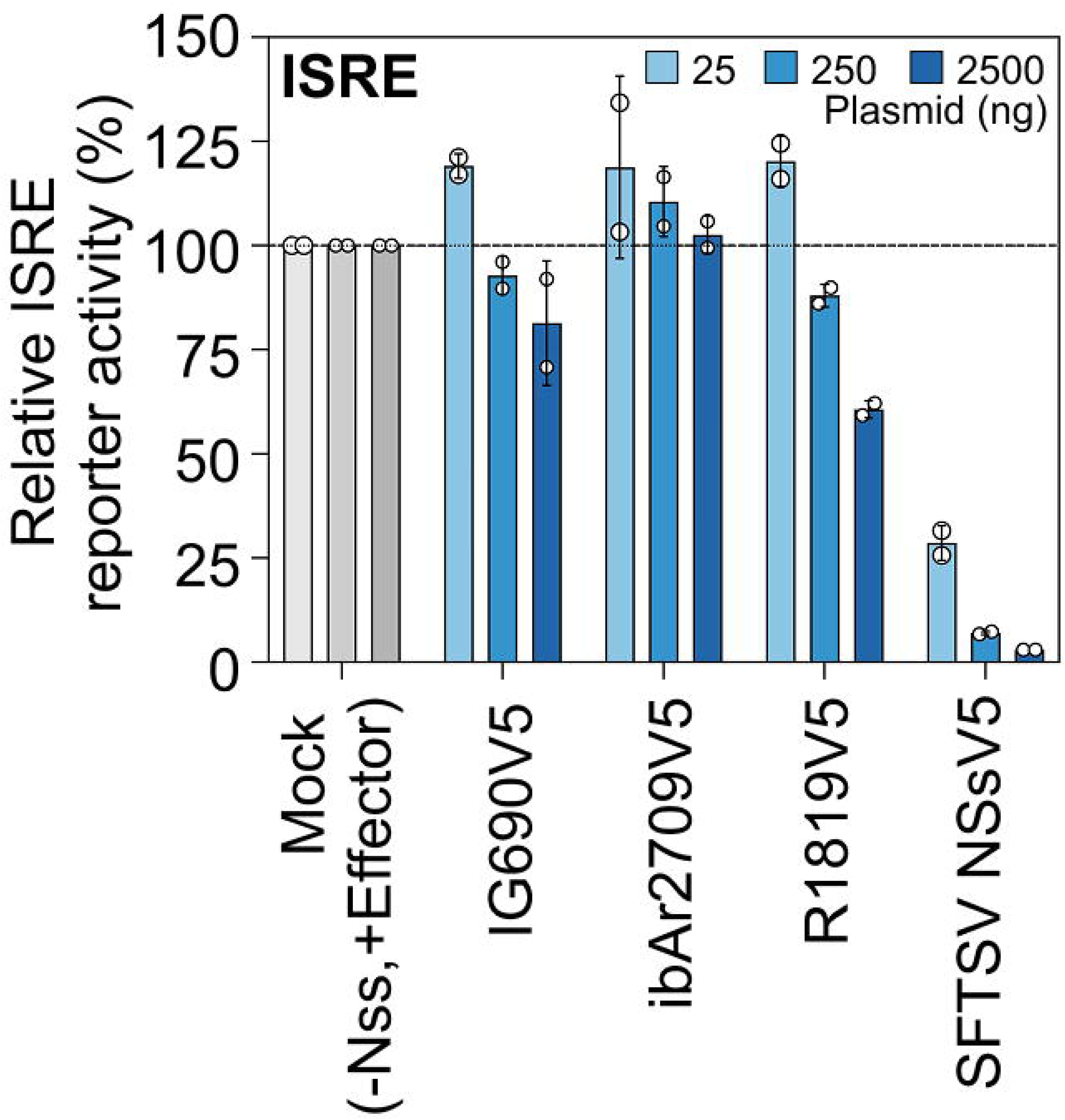
Bhanja virus NSs proteins weakly inhibit interferon signalling through the ISRE pathway. HEK-293T cells were co-transfected with an ISRE firefly luciferase reporter plasmid, pCMV-hRenilla for internal normalisation, and increasing amounts (25, 250 or 2500 ng) of plasmids expressing the indicated V5-tagged NSs proteins. SFTSV NSsV5 was included as a positive control for inhibition of interferon signalling. At 24 h post-transfection, cells were treated with IFNβ and lysed 24 h later for quantification of firefly and hRenilla luciferase activities. Firefly luciferase values were normalised to hRenilla and expressed as a percentage relative to IFNβ-treated cells lacking NSsV5 expression, which were set to 100% induction of the ISRE promoter (dashed line). The plotted values for each condition represent technical duplicates, with error bars indicating the range around the mean. Data are derived from one biological replicate.

Collectively, these results indicate that despite their capacity to inhibit interferon induction, BHAV NSs proteins are comparatively weak antagonists of downstream interferon signalling.

### Divergent NSs proteins modulate viral replication *in vivo*

An IFNAR-deficient mouse model was employed to assess the impact of NSs variation on viral pathogenesis. The IFNAR-deficient mouse model permits investigation of viral replication and NSs-independent innate immune restriction mechanisms. While this model does not permit assessment of NSs antagonism of canonical type I interferon signalling owing to the absence of IFNAR, it nevertheless retains intact upstream antiviral sensing pathways, including RIG-I-like receptor, Toll-like receptor and TBK1/IRF3 signalling cascades [60]. Groups of six IFNAR-deficient mice were infected subcutaneously with 10^4^ FFU of recombinant viruses and monitored for 7 days. All infected groups exhibited a similar pattern of weight change over the course of infection, with no clear differences in overall disease trajectory compared to each other, and values remaining largely within the range observed in mock-infected animals (Fig. 9A). Similarly, survival and clinical scoring revealed only mild disease across all groups, with no overt differences between viruses (SI Appendix, S7 Fig.).

**Fig 9.**
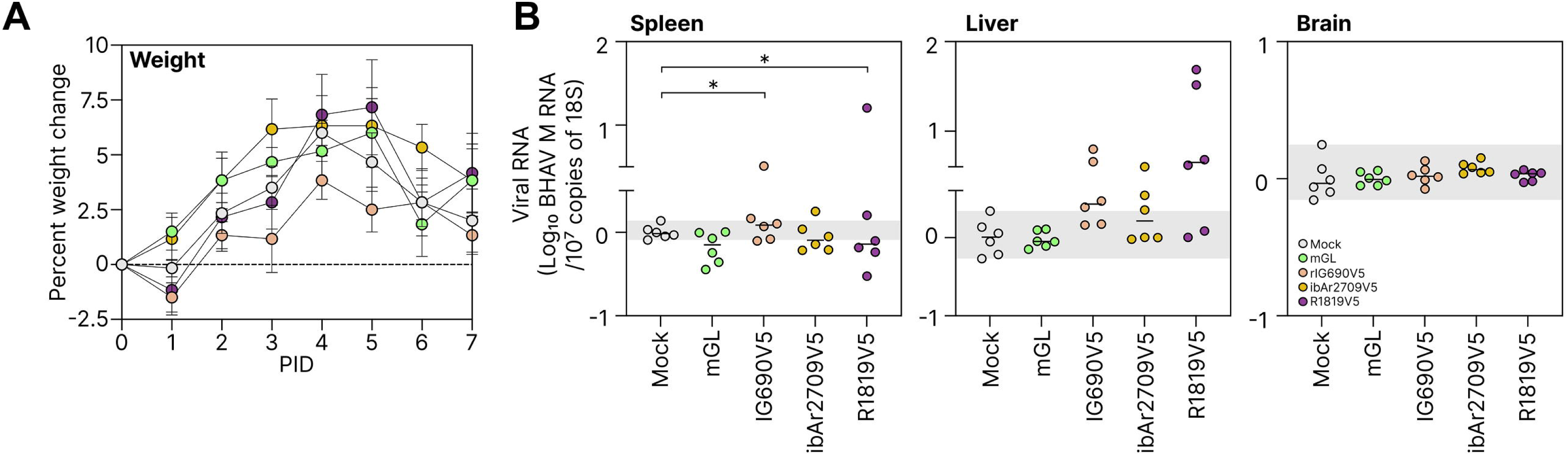
Divergent Bhanja virus (BHAV) NSs proteins modulate viral replication *in vivo*. **(A)** Percentage weight change of IFNAR-deficient mice (n=6 mice/group) following infection with recombinant BHAV expressing different NSs proteins. Groups of mice were mock infected (grey) or infected subcutaneously with 1 ×10^4^ PFU of rIG690 expressing mGreenLantern in place of NSs (rIG690ΔNSs-mGL; green), rIG690 expressing IG690 NSsV5 (orange), rIG690 expressing ibAr2709 NSsV5 (yellow) or rIG690 expressing R1819 NSsV5 (purple). Animals were monitored daily for changes in body weight over a 7-day period. Data represent the mean ± SD. The grey shaded region indicates the range of values observed in mock-infected animals. **(B)** Viral RNA levels in spleen, liver and brain tissues collected at 7 dpi from infected IFNAR-deficient mice. Viral RNA (M segment) was quantified by RT-qPCR and normalised to 18S RNA. Data are presented as Log_10_ BHAV RNA/18S RNA. Each point represents an individual animal, and horizontal bars indicate the mean. The grey shaded region indicates the range of values observed in mock-infected animals. Statistical significance was determined using one-way ANOVA with multiple comparisons. *P < 0.05.

Viral RNA levels were quantified in spleen, liver and brain at the experimental endpoint. In the spleen, animals infected with viruses expressing heterologous NSs proteins, particularly R1819NSsV5, showed increased viral RNA levels compared to the mGL control, with statistically significant differences observed (One-Way ANOVA, p= 0.0383) (Fig. 9B). A similar trend was observed in the liver, although variability between animals was greater and differences were less pronounced. In contrast, viral RNA levels in the brain were uniformly low across all infected groups and remained close to the limit of detection, indicating limited neuroinvasion under these conditions. Thus, NSs sequence variation can influence viral replication in peripheral tissues in vivo, particularly in the spleen.

## DISCUSSION

This study establishes the first reverse genetics platform for Bhanja virus and uses it to define how naturally occurring NSs sequence variation influences viral fitness and innate immune antagonism across divergent BHAV lineages. By combining virion RNA sequencing with terminal untranslated region mapping by 3′ RACE, we confirmed that our laboratory wtIG690 stock closely matched the previously published genome sequences while experimentally validating the viral segment termini required for recombinant virus recovery (Fig. 1). Using these validated sequences, we generated recombinant rIG690 entirely from cloned cDNA and demonstrated that it recapitulated the replication kinetics, viral protein expression and interferon sensitivity of the parental isolate in both interferon-competent and interferon-deficient cells (Fig. 2). Additionally, these tools enabled manipulation of the NSs locus, including generation of recombinant NSs swap viruses and an NSs-deficient reporter virus, thereby providing a tractable platform for comparative studies of BHAV molecular biology and pathogenesis.

Consistent with previous observations for several tick-borne phenuiviruses, BHAV replication was influenced strongly by the innate immune status of the infected cell [42,61,62]. wtIG690 replicated most efficiently in interferon-deficient or interferon-compromised cell systems, including Vero E6, Huh7/Huh7.5 and A549-VNPro cells, whereas replication remained substantially restricted in IFN-competent A549 cells (Fig. 3). This difference was reflected both in viral titres and N protein accumulation, supporting the conclusion that intact antiviral signalling constitutes a major barrier to productive BHAV replication in mammalian cells. Although BHAV has historically been associated with neurotropic disease in experimental animal infections, replication in the neuronal-derived cell lines examined here remained comparatively limited. This does not exclude productive infection of other neural cell populations, including astrocytes, glial cells or primary neuronal cultures. Together, these findings indicate that BHAV exhibits a relatively restricted mammalian cell tropism that is strongly shaped by host innate immune competence.

A similarly restricted phenotype was observed in arthropod-derived cell lines. Recombinant rIG690 replication was sustained in *R. microplus*-derived cells, whereas viral titres declined progressively in *I. ricinus*-, *I. scapularis*- and mosquito-derived cell cultures, accompanied by reduced or undetectable N protein accumulation by both western blotting and immunofluorescence (SI Appendix, S2 Fig.). These observations are notable given that historically, BHAV has been isolated from several ixodid tick genera, including *Haemaphysalis*, *Hyalomma*, *Dermacentor* and *Rhipicephalus* [2–4,63], yet has not been associated with *Ixodes* spp. ticks. The inability of rIG690 to maintain replication in *Ixodes*-derived cells may therefore reflect biologically relevant vector restrictions rather than a general incompatibility with arthropod systems. The preferential replication of rIG690 in *Rhipicephalus*-derived cells compared to *Ixodes*-derived cells reveals intriguing vector-specific determinants of BHAV permissiveness. This differential tropism may reflect species-specific differences in viral receptor expression, intracellular replication machinery, or constitutive antiviral response status [64,65]. Future studies employing cross-species transfection of candidate entry receptors and comparative transcriptomics of tick-derived cell lines will clarify the molecular basis of arthropod-specific BHAV tropism and its relationship to natural transmission cycles.

Similarly, the absence of productive replication in mosquito-derived cells is consistent with the lack of any recognised mosquito association for BHAV. Although these experiments do not directly establish vector competence, they support the concept that BHAV exhibits a comparatively restricted arthropod species tropism relative to more broadly permissive arboviruses [66].

One of the most striking findings of this study was the substantial divergence observed within the BHAV NSs protein despite the relative conservation of other viral proteins, particularly N [28]. To investigate the functional significance of this variation, recombinant rIG690 viruses expressing heterologous V5-tagged NSs proteins derived from African and European isolates were generated (Fig. 4). Viruses expressing ibAr2709NSsV5 or R1819NSsV5 exhibited enhanced viral protein accumulation and moderately increased replication in interferon-competent A549 cells relative to the parental IG690 backbone, whereas these differences were substantially reduced in A549-VNPro cells (Fig. 4). In contrast, IG690NSsV5 reproducibly exhibited lower expression levels across multiple experimental systems. Importantly, reduced activity of IG690 NSs was also observed using non-tagged constructs, indicating that these phenotypes were not solely attributable to epitope tagging. Notably, these differences were more apparent at the level of viral protein accumulation than endpoint viral titres. Although viruses expressing ibAr2709NSsV5 and R1819NSsV5 reproducibly exhibited increased N and NSs accumulation in interferon-competent cells, the corresponding increases in infectious virus production were comparatively modest (Fig. 4). This suggests that viral protein abundance may represent a more sensitive indicator of NSs-mediated fitness differences than peak virus titres under the experimental conditions employed. Together, these findings suggest that naturally-occurring NSs sequence variation contributes directly to differences in protein stability and/or innate immune antagonistic capacity between BHAV lineages.

Together, these findings indicate that BHAV remains highly sensitive to interferon-mediated restriction despite measurable differences in NSs antagonistic activity between isolates (Fig. 5). Although African- and European-derived NSs proteins suppressed IFN induction more efficiently than the prototype IG690 NSs, this did not translate into substantial resistance to exogenous IFNβ treatment. These observations suggest that BHAV NSs primarily functions to delay or dampen early antiviral sensing rather than to fully suppress establishment of the downstream interferon-induced antiviral state. Such a phenotype contrasts with highly pathogenic bandaviruses such as SFTSV, whose NSs proteins potently antagonise multiple stages of the interferon response [41,67,68]. Instead, the comparatively limited IFN antagonism exhibited by BHAV is consistent with its restricted replication in immune-competent mammalian cells and its overall attenuated phenotype.

Collectively, the data presented in Fig. 6 and Fig. 7 support a model in which BHAV NSs proteins primarily antagonise innate immune signalling at the level of TBK1/IKKε. Unlike SFTSV NSs, which potently sequesters multiple signalling components into prominent cytoplasmic inclusion bodies, BHAV NSs displayed a diffuse cytoplasmic localisation pattern and comparatively modest inhibitory activity. Nevertheless, the stronger antagonistic phenotypes associated with ibAr2709NSsV5 and R1819NSsV5 correlated with reduced TBK1 phosphorylation and increased spatial association with TBK1, suggesting that BHAV NSs may interfere with kinase activation through transient or proximity-dependent mechanisms rather than stable sequestration complexes. In this regard, BHAV NSs appears mechanistically more similar to HRTV NSs than to the canonical inclusion body-forming NSs proteins described for SFTSV and related bandaviruses [42,51]. The inability of the IG690-derived anti-NSs antibody to robustly detect heterologous NSs proteins is consistent with the marked sequence divergence observed within the NSs coding region.

In contrast to their effects on IFN induction, BHAV NSs proteins exhibited comparatively weak inhibition of IFN signalling through the ISRE pathway (Fig. 8). Even at high plasmid concentrations, BHAV NSs proteins only modestly reduced ISRE activation relative to the potent suppressive activity of SFTSV NSs. Together with the marked sensitivity of BHAV replication to exogenous IFNβ, these findings support a model in which BHAV NSs primarily acts to dampen early antiviral sensing pathways while exerting limited control over downstream interferon signalling. This comparatively restricted antagonistic profile may be relevant to the distinctive pathogenesis of BHAV. Although BHAV replicates poorly in many peripheral cell types and causes limited disease following peripheral inoculation in adult animals, severe neurological disease can occur following intranasal or intracranial challenge and in younger animals [16–20]. Whether this reflects adaptation to replication within specific cellular niches of the central nervous system remains unknown. Notably, the overall antagonistic capacity of BHAV NSs was substantially weaker than that reported for SFTSV NSs, which suppresses multiple stages of the innate immune response through formation of prominent cytoplasmic inclusion bodies that sequester key signalling molecules including TBK1, IRF3 and STAT proteins [37–41]. Although additional viral and host determinants undoubtedly contribute to disease outcome, the reduced ability of BHAV NSs to counteract innate immune responses may represent one factor limiting the pathogenic potential of BHAV relative to highly pathogenic bandaviruses such as SFTSV.

The biological relevance of these differences was further supported *in vivo* using an IFNAR-deficient mouse model (Fig. 9 and SI Appendix, S7 Fig.). Although all recombinant viruses caused only mild clinical disease over the seven-day infection period, viruses expressing an NSs protein generally accumulated to higher levels in peripheral tissues than the NSs-deficient control virus, with significant increases in splenic viral RNA observed for rIG690NSsV5 and rR1819NSsV5. Similar trends were evident in the liver. In contrast, viral RNA remained low in the brain across all groups, indicating limited neuroinvasion under these experimental conditions. Despite the absence of intact type-I interferon signalling in this model, these findings demonstrate that NSs sequence variation influences viral fitness and replication in peripheral tissues *in vivo*. The higher viral burdens observed in animals infected with viruses expressing the ibAr2709 and R1819 NSs proteins closely mirrored the greater ability of these NSs variants to suppress IFN induction pathways in vitro, supporting a functional link between NSs-mediated innate immune antagonism and enhanced viral replication in vivo. While the overall disease phenotype remained modest relative to highly pathogenic bandaviruses such as SFTSV [60], these data nevertheless establish NSs as an important determinant of BHAV replication and in vivo fitness.

Collectively, this study expands the currently limited molecular understanding of BHAV substantially and establishes a foundation for future investigations into the biology of tick-borne bandaviruses. The development of an authentic reverse genetics system, together with recombinant NSs-swap viruses and an NSs-deficient reporter virus, now enables direct investigation of viral determinants governing replication, immune evasion and host adaptation. More broadly, these findings demonstrate that relatively modest NSs sequence divergence can substantially alter innate immune antagonistic capacity and viral fitness while preserving an overall attenuated disease phenotype. Given the broad geographic distribution of BHAV and the limited surveillance currently performed for this virus, continued investigation into its ecology, vector associations and pathogenic potential is warranted.

## MATERIALS AND METHODS

### Experimental Model and subject detail

#### Vertebrate cell lines

Human cells: A549 (lung epithelial) [69], A549-NPro (expressing BVDV NPro; IRF-3 degradation) [54,70], A549-VNPro (generated by simultaneously and constitutively expressing PIV5-V and BVDV NPro; STAT1 and IRF-3 degradation) [71], Huh7.5-T7, Huh7 Lunet-T7, Huh7-ViperinKD (hepatocellular carcinoma; T7 polymerase-expressing or Viperin-knockdown variants) [72,73], SK-N-SH (neuroblastoma) [74], DAOY (medulloblastoma) [75], U373-MG (glioblastoma) [76], SVG p12 (astroglia, ATCC CRL-8621) [77]. Animal-derived cells: Vero E6 (African green monkey kidney) [69], BHK-21 (hamster kidney) [78], BSR-T7/5 (BSR expressing T7 polymerase) [79], MDBK (bovine kidney, ATCC CCL-22) [80], MDCK (canine kidney, ATCC CCL-34) [81], RAW264.7 (mouse monocyte/macrophage, ATCC TIB-71), MRK101 (vole kidney) [82], QT35 (Japanese quail fibrosarcoma, ECACC 93120832) [83], DF-1 (chicken fibroblast, ATCC CRL-12203) [84], E. Derm (horse fibroblast, ATCC CCL-57) [85].

Media: DMEM (ThermoFisher) with 10% heat-inactivated foetal bovine serum (FBS) for A549, Vero E6, BHK-21, BSR variants, and MDBK. EMEM (ThermoFisher) with 10% FBS and 1 mM sodium pyruvate for neuronal lines (SK-N-SH, DAOY, U373-MG). Huh7 variants and U373-MG supplemented with 1X non-essential amino acids (NEAA) (ThermoFisher). GMEM (ThermoFisher) with 10% FBS and 10% tryptose phosphate broth (TPB) for BHK-21 and BSR-T7/5. Selection maintained with 1 mg/ml G-418 (Promega; T7-expressing lines), 10 µg/ml Blasticidin (InvivoGen; A549-NPro, Huh7 variants), or 10 µg/ml Blasticidin + 2 µg/ml Puromycin (A549-VNPro). All cultured at 37°C, 5% CO₂.

#### Invertebrate cell lines

Tick-derived cells: IRE/CTVM19, IRE/CTVM20 [86], BME/CTVM6 [87], BME/CTVM23 [88], and ISE6 [89] sourced from the Tick Cell Biobank, University of Liverpool. Mosquito-derived cells: AF5 (*Ae. aegypti*) [90,91] and C6/36 (Aedes albopictus) [92].

Media: L-15 (Leibovitz) medium (Sigma-Aldrich) with 20% FBS, 10% TPB, and 2 mM L-glutamine for IRE/CTVM19, BME/CTVM6, and BME/CTVM23. L-15B medium [93] with 5% FBS, 10% TPB, 2 mM L-glutamine, 0.1% bovine lipoprotein cholesterol [MP Biomedicals]) for IRE/CTVM20 and ISE6; ISE6 additionally supplemented with 20% sterile water and pH neutralised with 1.0 N NaOH (Sigma-Aldrich). L-15 medium with 10% FBS, 10% TPB, 1X NEAA, 1 mM L-glutamine for AF5. DMEM with 10% FBS, 1X NEAA, 1 mM L-glutamine, 1X PenStrep (50 units/ml, ThermoFisher) for C6/36. Temperature: 28°C for IRE/CTVM19, IRE/CTVM20, BME/CTVM6, AF5, and C6/36; 32°C for BME/CTVM23 and ISE6. No CO₂ exposure.

#### Viruses

BHAV isolates: wtIG690 (1954, *Haemaphysalis intermedia*, Bhanjanagar, India) [1], wtibAr2709 (1964, *Rhipicephalus decoloratus*, Ibadan, Nigeria) [16], wtR1819 (1974, human laboratory worker, Colorado, USA infected by virus isolated from ticks in Yugoslavia) [22], wtR1329 (1974, *Haemaphysalis punctata*, Brač island, Croatia), and wtR1336 (1974, *Haemaphysalis punctata*, Brač island, Croatia) were provided by Brandy J. Russell (Arbovirus Reference Collection, Centers for Disease Control and Prevention, Fort Collins, USA).

All experiments involving wild-type and recombinant BHAV were performed under containment level 3 (CL-3) conditions, approved by the UK Health and Safety Executive.

#### Generation of virus working stocks

Wild-type and recombinant BHAV were propagated in A549-VNPro cells at MOI 0.01 FFU/cell. Infected cells were maintained at 37°C with 5% CO₂ until visible cytopathic effect appeared (typically days 7–10 p.i.). Supernatants were clarified (4000 xg, 5 min), aliquoted, and stored at −80°C. All viruses were used at passage 1, except rIG690ΔNSs:mGreenLantern (mGL) (passage 2).

#### Plasmids

Full-length antigenomic cDNA clones for BHAV IG690 S, M, and L segments were generated by RT-PCR from virion RNA and cloned into pTVT7 using In-Fusion Snap Assembly Master Mix (Takara Bio) [45,46,94,95]. The resulting plasmids (pTVT7-S, - M, -L) contained viral antigenomic sequences flanked by T7 promoter and hepatitis delta virus ribozyme. Expression constructs for BHAV N and RdRp/L were generated by cloning into pTM1 or pCMV backbones (pTM1-N/L and pCMV-N/L).

NSs swap viruses: C-terminal V5 epitope tag was introduced into pTVT7-S NSs open reading frame by site-directed mutagenesis. NSs coding sequences from ibAr2709, R1819, and R1329 were substituted into the IG690 S-segment backbone by In-Fusion cloning, generating pTVT7-S:ibAr2709NSsV5, pTVT7-S:R1819NSsV5, and pTVT7-S:R1329NSsV5. NSs-deficient control: NSs open reading frame replaced with mGreenLantern (mGL), generating pTVT7-SΔNSs:mGL. All plasmids propagated in JM109 (Promega) or C3040 (New England Biolabs) chemically competent *E. coli* (Zymo Research).

#### Production of BHAV IG690 anti-N and -NSs antibody

BHAV IG690 N and NSs coding sequences were inserted into modified pDEST14 vector with N-terminal hexahistidine tag (Invitrogen) [46,96]. Plasmids expressed in BL21(DE3) *E. coli* (New England Biolabs), induced with 1 mM IPTG, grown overnight at 20°C. Bacterial lysates prepared using B-PER reagent with protease inhibitor cocktail, lysozyme, and DNase I (ThermoFisher). N protein purified from soluble lysates via HisTrap High Performance column (Cytiva) with wash buffer (80 mM imidazole, 0.5 M NaCl, 50 mM Tris-HCl pH 8.0, 10% glycerol, 0.1% Triton X-100, 0.1% Tween-20, 0.1% DTT) and eluted with 500 mM imidazole in equivalent buffer. NSs protein solubilized from insoluble lysates and purified using ionic and denaturing buffers [97]. Both proteins buffer-exchanged into PBS (dialysis cassettes, 10K MWCO) and concentrated (5K MWCO). Protein concentration quantified using BSA standards; samples supplied to Eurogentec for rabbit immunization.

#### Virus Rescue

BSR-T7/5 cells (3×10^5^/well, 6-well plates) were seeded in DMEM + 10% FBS. Next day, medium was replaced with DMEM with 5% FBS and cells co-transfected with 400 ng transcription plasmids (pTVT7-S, -M, -L) and 200 ng helper plasmids (pCMV-N, -L). For NSs variants, pTVT7-S was replaced with pTVT7-S:NSsV5, pTVT7-S:ibAr2709NSsV5, pTVT7-S:R1819NSsV5, pTVT7-S:R1329NSsV5, or pTVT7-SΔNSs:mGL. After 6 days at 33°C, supernatants were clarified (4000 xg, 10 min) and used to propagate working stocks in A549-VNPro cells. Virus recovery assessed by focus-forming assay or mGreenLantern signal (EVOS M7000, Invitrogen).

#### Virus titration

Vero E6 cells were seeded in 12-well plates, infected next day with serial virus dilutions in DMEM containing 2% FBS. Inoculum was incubated 1 h at 37°C, followed by overlay (1X MEM supplemented with 0.6% Avicel and 2% FBS). After 6 days at 37°C, cells were fixed in 8% formaldehyde. For antibody detection, cells were permeabilized in PBS containing 0.5% Triton X-100 and 20 mM sucrose; probed overnight at 4°C with BHAV anti-N antibody (1:1000 in PBS-T containing 5% milk); washed in PBS-T; probed 2.5 h at room temperature with goat anti-rabbit HRP (1:2000, Sigma, in PBS-T containing 5% milk); washed in PBS-T; developed with SeraCare TrueBlue Peroxidase Substrate (Insight Biotechnology) for 15–20 min; final wash in distilled water to prevent substrate overexposure.

#### Virus Infection

Cells were seeded into 12- or 96-well plates. After overnight incubation, cells were inoculated with low-volume in DMEM 2% (v/v) FBS containing viruses that had been diluted to a desired MOI. After 1h incubation at 37°C, virus inoculum was removed and replenished with fresh cell culture medium and was considered the 0h p.i. timepoint. At desired times p.i., culture medium supernatants and cell monolayer lysates were harvested and analysed.

#### Mice

All mice were 8 to 12-week-old interferon alpha receptor knockout (IFNAR-/-) mice on a 129S7/SvEvBrdBkl-Hprt^b-m2^ background (Marshall Bioresources), comprising both male and female animals. A129 (IFNAR-/-) mice were purchased from Marshall Bioresources and maintained in a pathogen-free facility, in filter-topped cages, and maintained in accordance with local and governmental regulations. All mice had a 12-h light/12-h dark cycle and were provided with sterile food and tap water *ad libitum*.

### Infection of mice with viruses

Mice were anaesthetised by isoflurane inhalation and injected with 1×10^4^ foci forming units of BHAV in 100µl of DMEM, subcutaneously into the right flank using a 26G needle. Mice were monitored for moderate signs of infection (SI Appendix, S2 Table) and culled humanely when they reached a clinically defined humane endpoint of disease or at specified timepoints.

### RNA extraction from tissues

Tissue samples were homogenised in TRIzol (Life Technologies) using a Precellys 24 (Bertin instruments) with 7mm metal beads (Qiagen), followed by purification using PureLink columns with DNase digestion (Life Technologies).

### Gene expression analysis

Viral RNA and host gene transcripts were quantified by reverse transcription qPCR. Tissues (up to 100 mg) yielded total RNA; 1 ug RNA was converted to cDNA using High-Capacity RNA-to-cDNA kit (Life Technologies), then 1% used per qPCR assay (10 ng RNA equivalent). BHAV-specific qPCR primers targeted genomic M RNA. RNA was extracted using PureLink Plus columns. qPCR analysis was performed using SYBR-green (Applied Biosystems) on a QuantStudio 7 Flex Real-Time PCR System (Applied Biosystems). Primer details in SI Appendix, S1 Table.

### Virion RNA extraction and RT-PCR

Supernatant stocks containing BHAV underwent virion RNA extraction using the QIAamp Viral RNA Mini Kit (Qiagen). Reverse transcription (RT) of isolated RNA into complementary DNA (cDNA) was carried out using segment specific primers (SI Appendix, S1 Table) and SuperScript III Reverse Transcriptase (Invitrogen), as per the manufacturer’s instructions. Amplification of viral cDNA by Polymerase Chain Reaction (PCR) was carried using Q5 High-Fidelity DNA Polymerase (New England Biolabs), as per the manufacturer’s instructions. After analysis by agarose gel electrophoresis, PCR products were gel-purified with Wizard SV Gel and PCR Clean-Up System (Promega) and subject to Sanger sequencing (Eurofins Genomics or Source Bioscience).

### 3’ RACE

Virion RNA was isolated from cell-culture supernatants (QIAamp Viral RNA Mini Kit, Qiagen). Total cellular RNA was extracted from wtIG690-infected monolayers (TRIzol, Invitrogen). RNAs were polyadenylated (Poly(A) Tailing Kit, Invitrogen) and purified (RNeasy Mini Kit, Qiagen). Polyadenylated RNA (10 µl) was reverse transcribed to cDNA using GoScript (Promega) or SuperScript III (Invitrogen) reverse transcriptase with 100 µM Oligo(dT)15 primer (SI Appendix, S2 Table) at 42–50°C for 60 min. cDNA (5 µl) was PCR amplified with Q5 High-Fidelity DNA Polymerase (New England Biolabs) using RACE anchor primer and segment-specific primers (SI Appendix, S1 Table). PCR products were analysed by agarose gel electrophoresis, gel purified, and sequenced (Sanger) to determine nucleotide sequences.

### Western blot

Cell lysates were harvested in Laemmli lysis buffer (100 mM Tris-HCl pH 6.8, 4% SDS, 20% glycerol, 0.2% bromophenol blue, 200 mM DTT, 22.5U Pierce Universal Nuclease [ThermoFisher]), boiled 10 min at 95°C and stored at −80°C. Samples were separated on 4–12% NuPAGE Bis-Tris gels (Invitrogen) at 140–180 V for 1 h, transferred to 0.45 μm nitrocellulose membranes (Amersham) using Trans-Blot Turbo System (Bio-Rad) at 10 V for 50 min. Membranes were blocked in PBS-T (0.1% Tween-20, 5% milk) for 1 h and probed overnight at 4°C with BHAV anti-N and anti-NSs rabbit polyclonal antibodies, or commercial antibodies: anti-Tubulin (T5168, Sigma-Aldrich), anti-V5 tag (ab27671, Abcam), anti-TBK1/NAK (3013, Cell Signaling Technology), anti-phospho-TBK1/NAK Ser172 (5483, Cell Signaling Technology), anti-IR, F3 (9082, Cell Signaling Technology), anti-phospho-IRF3 [s386] (76493, Abcam), anti-FLAG (14793, Cell Signaling Technology), anti-Actin (A5060, Sigma Aldrich). After PBS-T washes, cells were incubated 1.5 h at room temperature with goat anti-mouse IgG H+L DyLight 800 or goat anti-rabbit IgG H+L DyLight 680 (Invitrogen), washed, and visualized on an Odyssey DLx Imaging System (LI-COR Biosciences).

### IFN**β** promoter reporter assays

HEK-293T cells were seeded into 24-well plates and co-transfected using TransIT-LT1 (Mirus Bio) with plasmids expressing BHAV, SFTSV or HRTV NSs proteins together with expression constructs encoding components of the IFN induction pathway (RIG-I, MDA5, MAVS, TBK1, IKKε or IRF3-5D, [98,99]). To measure IFNβ promoter activation, cells were co-transfected with the firefly luciferase reporter plasmid pIFN(-125)FfLuc [100] and pCMV-hRenilla as an internal transfection control. BHAV NSs expression plasmids were transfected at 25, 250 or 2500 ng per well, while IFN pathway effector plasmids were transfected at 50 ng per well. Total DNA input was equalised using empty pCMV vector. At 24 h post-transfection, cells were lysed and firefly and Renilla luciferase activities quantified using a dual-luciferase reporter assay. Firefly luciferase activity was normalised to Renilla luciferase activity and expressed as a percentage relative to cells transfected with the corresponding empty vector control. FLAG-tagged RIG-I N, MAVS, TBK1, IKKε, and IRF3-5D expression plasmids were kindly provided by Mirko Schmolke (University of Geneva) [36].

### ISRE reporter assays

HEK-293T cells were transfected as described above, except that the pIFN(-125)FfLuc reporter plasmid was replaced with p(9–27)4tkΔ(−39)lucter (referred to as pISRE-FFLuc, [101]), containing firefly luciferase under the control of interferon-stimulated response elements (ISREs). Twenty-four h post-transfection, cells were treated with recombinant human IFNβ (250 U/well) or mock treated. After a further 24 h incubation, cells were lysed and firefly and Renilla luciferase activities measured. Firefly luciferase activity was normalised to Renilla luciferase activity and expressed as a percentage relative to IFNβ-treated cells transfected with the corresponding empty vector control.

### CPE protection assay

IFN-competent A549 cells grown in 12-well plates were infected with viruses at MOI 2 FFU/cell. Supernatants were harvested at 24 h p.i. and inactivated by UV irradiation (8 W, 254 nm, 2 cm distance, 4 min with occasional shaking). A549 cell monolayers were lysed in Laemmli buffer for western blot analysis. Inactivated supernatants underwent two-fold serial dilution in fresh medium containing 2% FBS. Sub-confluent IFN-incompetent A549-NPro cell monolayers (96-well plates) were pre-treated with dilution series for 24 h, then infected with IFN-sensitive Encephalomyocarditis virus (EMCV) at MOI 0.05 PFU/cell. Monolayers were fixed in 8% formaldehyde at 4 days p.i., stained with 0.2% bromophenol blue to visualize EMCV cytopathic effect.

### Immunofluorescence

Tick and mosquito cells were seeded overnight in Ibidi 8-well high chamber µ-slides (Thistle Scientific) and were subsequently infected at MOI 1 FFU/cell. Cells were fixed at day 12 p.i. in 4% formaldehyde for 1.5 h, permeabilized for 30 min with PBS containing 0.5% Triton X-100 (Fisher Scientific) and 20 mM sucrose and probed overnight at 4°C with rabbit IG690 anti-N in PBS containing 2% FBS. After PBS-T washes, the cells were probed 1.5 h with goat anti-rabbit Alexa Fluor 594 (A-11037, Invitrogen) in PBS containing 2% FBS, mounted in Vectashield antifade mounting medium with DAPI (2BScientific) and imaged by confocal microscopy (Zeiss LSM 880).

### Bioinformatic analysis

Raw sequencing reads from Bhanja virus sample Bhanja_S1_L001 were processed using the bioinformatic pipeline: Forward reads (R1) were trimmed using fastp v0.23.4 with quality filtering (Q≥30, minimum read length 50 bp) and adapter sequence removal. Trimmed reads were aligned to the Bhanja virus strain IG690 reference genome (NCBI GenBank: JX961619.1, JX961620.1, JX961621.1; total 11,501 bp across 3 RNA segments) using Bowtie2 v2.5.2 with sensitive alignment mode (parameters: -D 15 -R 2 -L 22 -i S,1,1.15). Alignments were converted to BAM format, sorted by coordinate, and indexed using SAMtools v1.19.2. Per-nucleotide read depth was calculated using samtools depth across the entire reference genome.

### Statistical Analysis

Mouse survival curves were undertaken once on ethical grounds in line with NC3Rs policy and the ARRIVE Guidelines 2.0 (Percie du Sert N et al. (2020) Reporting animal research: Explanation and elaboration for the ARRIVE guidelines 2.0. PLOS Biology 18(7): e3000411. https://doi.org/10.1371/journal.pbio.3000411 (SI Appendix, S7A Fig). *In vivo* experiments were performed with appropriate animal numbers to achieve a 90% power with a significance level (alpha) of 0.05 (two-tailed), calculated with StatMate 2.0 (GraphPad).

All *in vitro* studies and microscopy were undertaken on a minimum of two separate occasions with three technical replicates to ensure reproducibility. All results were confirmed between experimental replicates.

Data were analysed using GraphPad Prism Version 11.0.1 software. Testing for statistical significance was carried out by either unpaired t-test, one-way ANOVA, or two-way ANOVA with Sidak’s multiple comparison. The cut-off for significance was set at p<0.05. Where analysis was employed, unless stated otherwise both “ns” and a lack of notation indicate no significance.

### Ethics statement

Animals were maintained at the University of Glasgow under specific pathogen-free conditions in accordance with UK Home Office regulations (Project License PP8216562), the Animals (Scientific Procedures) Act 1986, and University of Glasgow ethics committee approval. All animal research adhered to ARRIVE guidelines. Mice were humanely euthanized upon exhibiting three or more moderate clinical signs, >15% body weight loss, or at predefined endpoints. Clinical scoring: 0 = no signs, 1 = mild signs, 2 = moderate signs, 3 = one advanced or three moderate signs (see SI Appendix, Table S2).

## ACKNOWLEDGEMENTS

We thank Prof. Ulrike Munderloh, University of Minnesota, for permission to use the ISE6 cell line. We also thank Prof. Richard Randall and Dr Dan Young, University of St Andrews, for provision of, and permission to use, the A549-Npro and A549-VNPro cell lines. Finally, we thank Dr James Dunlop for advice on protein expression.

## FUNDING

This research was funded by the University of Glasgow MVLS DTP (A.T.C) and a Wellcome Trust/Royal Society Sir Henry Dale Fellowship (210462/Z/18/Z) (B.B.). The facilities used in this study were funded by the UK Medical Research Council (MC_UU_00034/8, MC_UU_00034/7). L.B.S. was supported by the Wellcome Trust grant no. 223743/Z/21/Z. A.K. was funded by the UK Medical Research Council (MC_UU_12014/8, MC_UU_00034/4, MC_UU_00034/5/RCUK).

The authors acknowledge that the *in vivo* mouse work was carried out by CTH: CVR Translational Hub at the MRC-University of Glasgow Centre for Virus Research (CVR), funding for which in part was supported by MRC and LifeArc.

This research was funded in whole or in part by the Wellcome Trust. The funders had no role in study design, data collection and analysis, decision to publish, or preparation of the manuscript.

## AUTHORS’ CONTRIBUTIONS

Conceptualization: A.T.C., A.K., B.B., Data Curation: B.B., Formal Analysis: A.T.C., A.K., B.B., Funding Acquisition: B.B., Investigation: A.T.C., K.D., M.F., A.S., D.C., K.K., V.H., Methodology: A.T.C., K.D., M.F., A.S., D.C., K.K., V.H., B.B., Project administration: B.B., Resources: L.B.S., A.H.P, B.B., Software: Supervision: M.F., D.C., B.J.W., A.H.P, A.K., B.B., Validation: A.T.C., A.K., B.B., Visualization: A.T.C., B.B., Writing – original draft: A.T.C., B.B., Writing – review & editing: A.T.C., K.D., M.F., D.C., K.K., V.H., L.B.S., B.J.W., A.H.P, A.K., B.B.

## DECLARATION OF INTERESTS

The authors declare no competing interests.

## DATA AVAILABILITY

The data that support the findings of this study are openly available from Enlighten Research Data (https://doi.org/10.5525/gla.researchdata.2291). All sequencing data generated in this study have been deposited in the NCBI Sequence Read Archive under BioProject accession number PRJNA1476151. The NSs coding sequences of BHAV isolates R-1329 and R-1336 were deposited in GenBank under accession numbers PZ464570 and PZ464571, respectively.

## Notes

### Competing Interest Statement

The authors have declared no competing interest.

