## Supplementary Information for "Functional divergence of the NSs protein defines interferon antagonism and viral fitness across Bhanja virus lineages"

**The PDF file includes:**

S1 Fig. Mapping of transcription termination sites within the Bhanja virus wtIG690 S segment by 3′ RACE.

S2 Fig. BHAV rIG690 is not maintained in arthropod-derived cell lines.

S3 Fig. Multiple sequence alignment of BHAV isolate NSs proteins.

S4 Fig. Differential expression and proteasomal sensitivity of V5-tagged BHAV NSs proteins.

S5 Fig. Sensitivity of recombinant BHAV expressing heterologous V5-tagged NSs proteins to IFNβ treatment.

S6 Fig. Subcellular localisation of V5-tagged NSs proteins from BHAV and SFTSV.

S7 Fig. Clinical assessment of IFNAR-deficient mice infected with recombinant BHAV expressing heterologous NSs proteins.

S1 Table. Oligonucleotides used in this study.

S2 Table. Scoring system for the welfare assessment of virus challenged experimental mice.

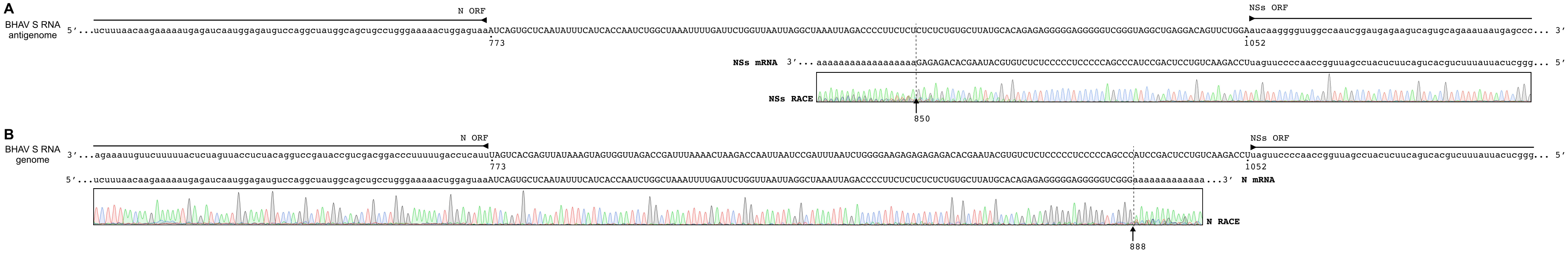


**S1 Fig. Mapping of transcription termination sites within the Bhanja virus wtIG690 S segment by 3′ RACE.**

**(A)** Determination of the NSs mRNA transcription termination site within the wtIG690 S segment. RNA extracted from wtIG690-infected Vero E6 cells was subjected to 3′ RACE using primers specific for the NSs ORF. Representative Sanger sequencing chromatograms are shown. The NSs mRNA transcription termination site within the S segment intergenic region is indicated at nucleotide position 850.

**(B)** Determination of the N mRNA transcription termination site within the wtIG690 S segment. RNA extracted from wtIG690-infected Vero E6 cells was subjected to 3′ RACE using primers specific for the N ORF. Representative Sanger sequencing chromatograms are shown. The N mRNA transcription termination site within the S segment intergenic region is indicated at nucleotide position 888.

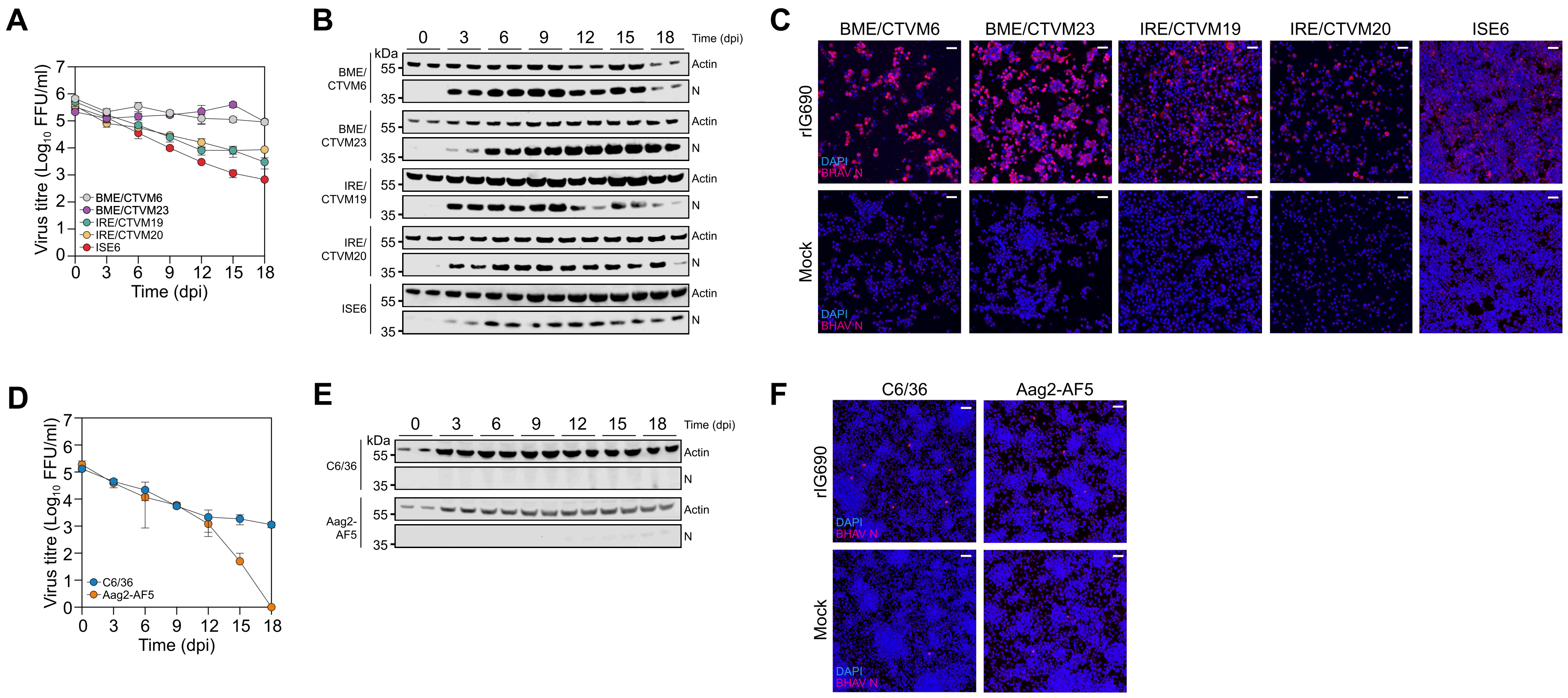


**S2 Fig. Growth kinetics of BHAV rIG690 in arthropod-derived cell lines.**

**(A)** Growth kinetics of recombinant BHAV rIG690 in tick-derived cell lines. BME/CTVM6 and BME/CTVM23 (*Rhipicephalus microplus*), IRE/CTVM19 and IRE/CTVM20 (*Ixodes ricinus*) and ISE6 (*Ixodes scapularis*) cells were infected with BHAV rIG690 at an MOI of 1 FFU/cell and culture supernatants were harvested at the indicated time points for titration by focus-forming assay on Vero E6 cells. Viral titres are presented as $Log_{10}$ FFU/ml. Data represent the mean ± SD from three biological replicates.

**(B)** Detection of BHAV infection in tick-derived cell lines by western blotting. Cell lysates collected at the indicated time points p.i. were analysed using BHAV anti-N and anti-Actin antibodies. Representative blots are shown.

**(C)** Anti-N immunofluorescent staining of tick-derived cell lines infected with BHAV rIG690. Tick-derived cell lines were either mock infected or infected with BHAV rIG690 at an MOI of 1 FFU/cell. Cells were fixed at 12 d p.i. and stained using a BHAV anti-N antibody (red) and DAPI (blue). Images were acquired using a Zeiss LSM 880 confocal microscope. Scale bar, 50 µm.

**(D)** Growth kinetics of recombinant BHAV rIG690 in mosquito-derived cell lines. C6/36 (*Aedes albopictus*) and Aag2-AF5 (*Aedes aegypti*) cells were infected with BHAV rIG690 at an MOI of 1 FFU/cell and culture supernatants were harvested at the indicated time points for titration by focus-forming assay on Vero E6 cells. Viral titres are presented as Log10 FFU/ml. Data represent the mean ± SD from three biological replicates.

**(E)** Detection of BHAV infection in mosquito-derived cell lines by western blotting. Cell lysates collected at the indicated time points p.i. were analysed using BHAV anti-N and anti-Actin antibodies. Representative blots are shown.

**(F)** Anti-N immunofluorescent staining of mosquito-derived cell lines infected with BHAV rIG690. Mosquito-derived cell lines were either mock infected or infected with BHAV rIG690 at an MOI of 1 FFU/cell. Cells were fixed at 12 d p.i. and stained using a BHAV anti-N antibody (red) and DAPI (blue). Images were acquired using a Zeiss LSM 880 confocal microscope. Scale bar, 50 µm.

```
IG690        MDDSRSIQFRLSGEVSPGSLIVGGYWRMSVANVRIRDIHTLYPPVPRELVDLFQGSAAIQ    60
ibAr2709     MDDSRSIQFRLSGEVSPGSLIIGGYWRMSVANIRIRDVHTLYPPVPRELADLFQGTAAIQ    60
R-1329       MDDSRSIQFRLSGEVSPGSLIIGGYWRMSVANIRIRDVHTLYPPVPRELVDLFQGTAAIQ    60
R-1336       MDDSRSIQFRLSGEVSPGSLIIGGYWRMSVANIRIRDVHTLYPPVPRELVDLFQGTAAIQ    60
R-1819       MDDSRSIQFRLSGEVSPGSLIIGGYWRMSVANIRIRDVHTLYPPVPRELVDLFQGTAAIQ    60
             *********************:**********:****:***********.*****:****

IG690        TLFLDGGHMVRSLPRPIFLFETMEDILPKGRPMTSQERSSLRWPQGVASISWISLALKVY    120
ibAr2709     TLFLDGGHMVRSLPRPLYLLGTMEEILPKGRPMTKQERSSLRWPQNVASVSWISLALKVY    120
R-1329       TLFLDGGHMVRSLPRPIFLLETMEDILPKGRPMTSQERSSLRWPQGVTSVSWISLALKVY    120
R-1336       TLFLDGGHMVRSLPRPIFLLETMEDILPKGRPMTSQERSSLRWPQGVTSVSWISLALKVY    120
R-1819       TLFLDGGHMVRSLPRPIFLLETMEDILPKGRPMTSQERSSLRWPQGVTSVSWISLALKVY    120
             ****************::*: ***:*********.**********.*:*:**********

IG690        GNGTPLERCFSWKALVRLLIRSTPESDPESSVADRLYDLNQKANKRAQRMGLMSALGGSP    180
ibAr2709     GNGTPLERCFSWKALVKLLIRSTPESCRESAIADRLYDLDQKANIRAQRLGVFSALSGTP    180
R-1329       GHGTPLERCYSWKALVKLLIKSTPESNHESTIADRLFDLNRKANVRAQRLGVLGVLSGCP    180
R-1336       GHGTPLERCYSWKALVKLLIKSTPESNHESTIADRLFDLDRKANVRAQRLGVLGVLSGCP    180
R-1819       GHGTPLERCYSWKALVKLLIKSTPESNHESTIADRLFDLNRKANVRAQRLGVLGVLSGCP    180
             *:*******:******:***:*****  **::****:**::*** ****:*::..*.* *

IG690        ILMKVGLLQALSYLRAAAKDLELLGITTSGLVDVQVLQALGWYYPVKLPRPIRKQVDEYI    240
ibAr2709     ILMKVGFLQALSYLRMAAKDLELLGISNSGLVDVQVLQALSWFYPVKLPRPIRKSVDDYI    240
R-1329       ILMKVGLLQVLSYLRAAAKDLELLGISTSGLVDVQVLQALGWFYPVKLPRPIRKEVDEYI    240
R-1336       ILMKVGLLQVLSYLRAAAKDLELLGISTSGLVDVQVLQALGWFYPVKLPRPIRKEVDEYI    240
R-1819       ILMKVGLLQVLSYLRAAAKDLELLGISTSGLVDVQVLQALGWFYPVKLPRPIRKEVDEYI    240
             ******:**.***** **********:.************.*:***********.**:**

IG690        RIEASQGSNTRILLLETFESGDRRKINRGPPFNEADFLSGLSCSESIEYFLETQESFRAH    300
ibAr2709     RIEASLGSNTRILLLETFDSRDRKKVNKGPPFDEATHLSKLSCEESINYFLETQETFRLH    300
R-1329       RIETSQGSNSRILLLETFSSGDRRKVNRGPPFDEASHLAKLSCDESITYFLETQETFRLH    300
R-1336       RIEASQGSNTRILLLETFNSGDRRKVNRGPPFDEASHLAKLSCDESITYFLETQETFRLH    300
R-1819       RIEASQGSNTRILLLETFNSGDRRKVNRGPPFDEASHLAKLSCDESITYFLETQETFRLH    300
             ***:* ***:********.* **:*:*:****:** .*: ***.*** *******:** *

IG690        YFCTDFSSDWPTP    313
ibAr2709     YFCTKFTSDWPTP    313
R-1329       YFCTEFTSDWPTP    313
R-1336       YFCTQFTSDWPTP    313
R-1819       YFCTQFTSDWPTP    313
             ****.*:******
```

**S3 Fig. Multiple sequence alignment of BHAV isolate NSs proteins.**

Clustal alignment of NSs sequences from five Bhanja virus isolates (IG690, ibAr2709, R-1329, R-1336, and R-1819). Amino acid residues are coloured by chemical properties (red, acidic/basic; blue, hydrophobic; green, polar). Asterisks (*), colons (:), and periods (.) indicate identical, strongly similar, and weakly similar residues, respectively.

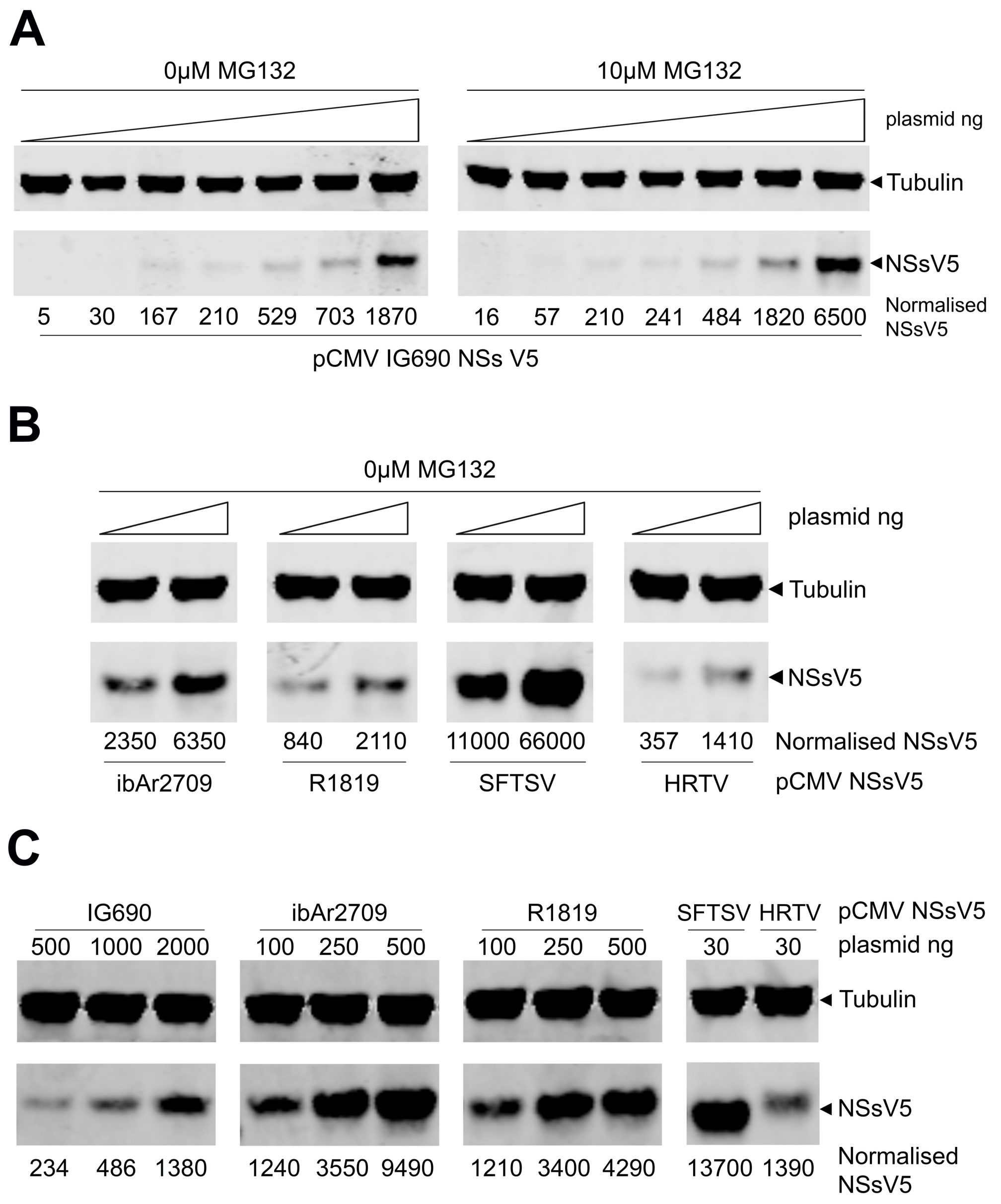


**S4 Fig. Differential expression and proteasomal sensitivity of V5-tagged BHAV NSs proteins.**
**(A)** HEK-293T cells were transfected with increasing amounts (0, 10, 50, 100, 250, 500 or 1000 ng) of a plasmid expressing V5-tagged IG690 NSs from a pCMV backbone. At 24 h post-transfection, cell lysates were harvested and analysed by western blotting using anti-V5 and anti-tubulin antibodies.
**(B)** HEK-293T cells were transfected with plasmids expressing V5-tagged NSs proteins from BHAV isolates ibAr2709 and R1819, or from SFTSV and HRTV, at either 250 ng or 500 ng. At 24 h post-transfection, cells were treated with either the proteasome inhibitor MG132 (10 μM) or DMSO control for 4 h prior to lysis. Cell lysates were analysed by western blotting using anti-V5 and anti-tubulin antibodies. NSsV5 signal intensities were normalised to tubulin.
**(C)** HEK-293T cells were transfected with varying amounts of plasmids expressing V5-tagged BHAV NSs proteins (500, 1000 or 2000 ng), or lower amounts of SFTSV NSsV5 (30 ng) or HRTV NSsV5 (30 ng), to achieve broadly comparable protein expression levels. At 24 h post-transfection, cell lysates were harvested and analysed by western blotting using anti-V5 and anti-tubulin antibodies. V5 signal intensities were internally normalised to tubulin.

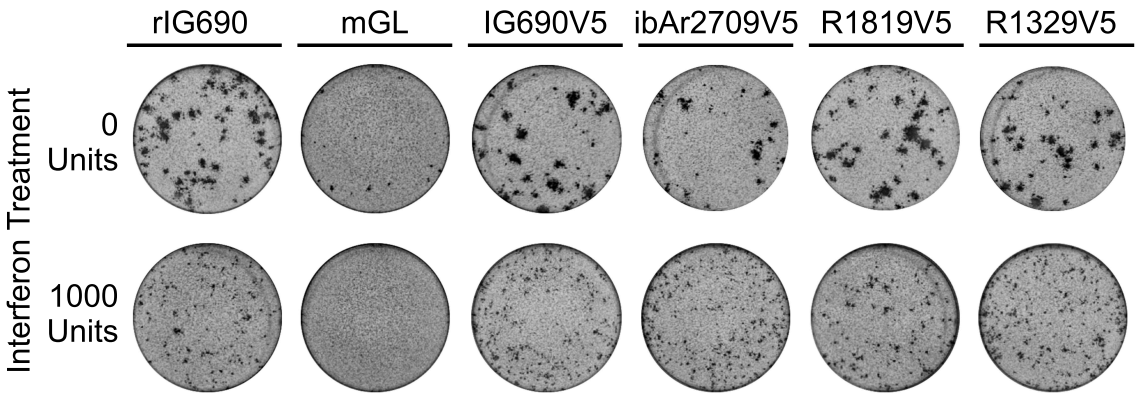


**S5 Fig. Sensitivity of recombinant BHAV expressing heterologous V5-tagged NSs proteins to IFNβ treatment.**

Representative anti-N immunostained monolayers from the immunofocus assays described in Fig. 5B. Images were acquired at dilutions permitting clear visualisation of individual foci to compare focus morphology and size between recombinant viruses following IFNβ treatment.

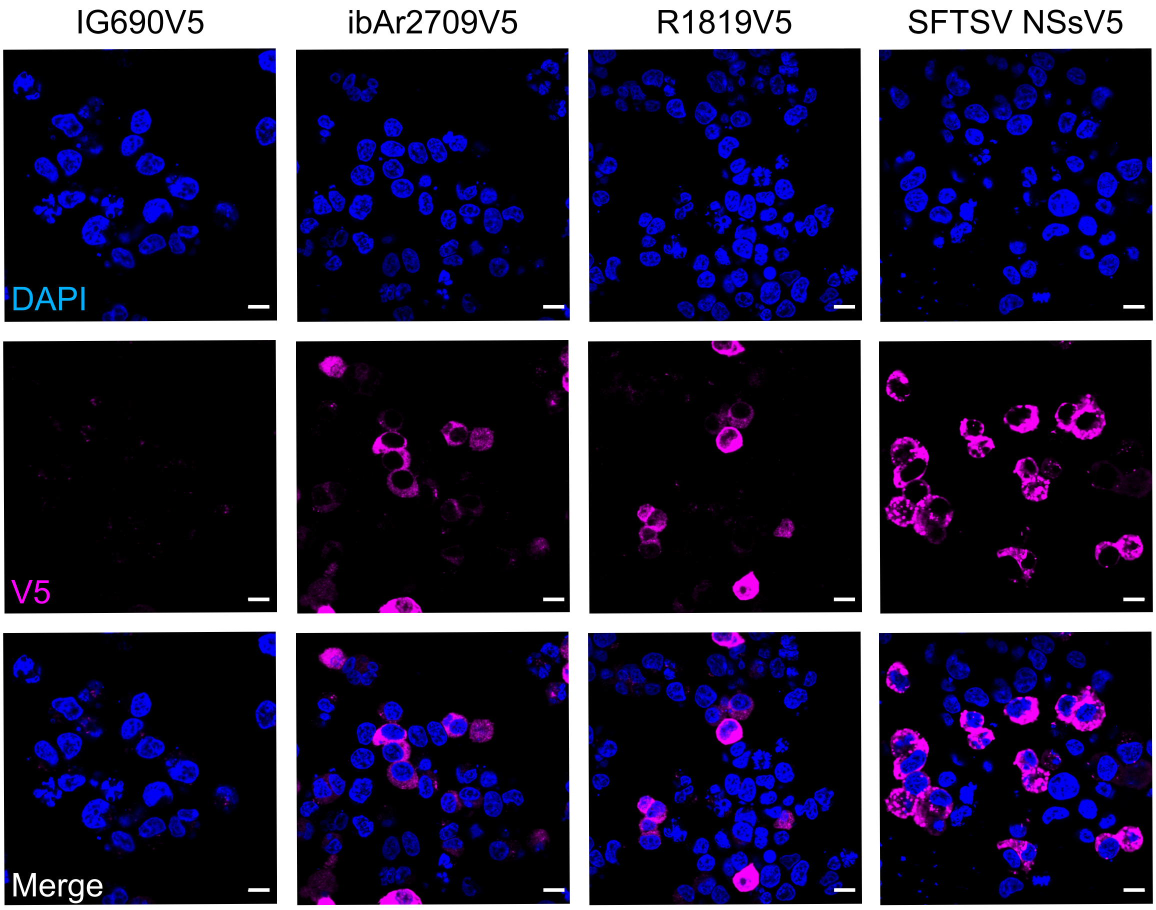


**S6 Fig. Subcellular localisation of V5-tagged NSs proteins from BHAV and SFTSV.**
HEK-293T cells were transfected with 500 ng of plasmids expressing the indicated C-terminally V5-tagged NSs proteins. At 24 h post-transfection, cells were fixed, permeabilised and stained using an anti-V5 antibody. Nuclei were counterstained with DAPI. Images were acquired using a Zeiss LSM 710 confocal microscope. Scale bars, 10 μm.

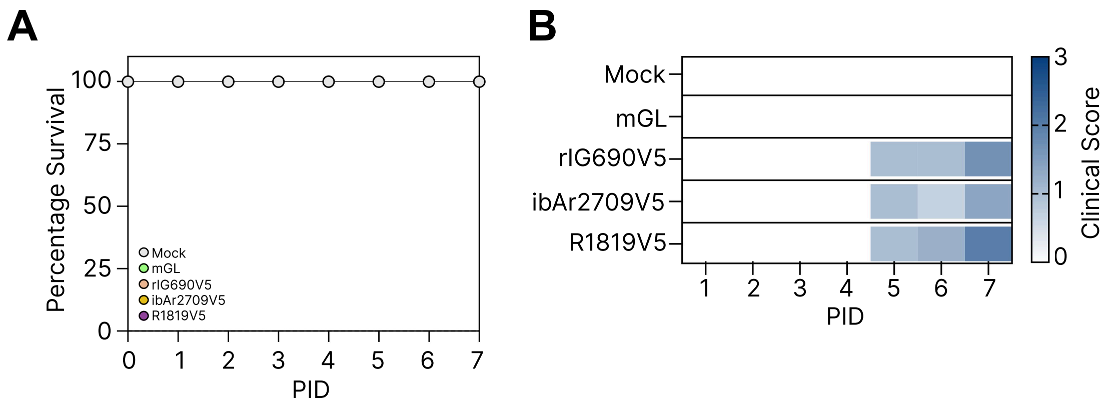


**S7 Fig. Clinical assessment of IFNAR-deficient mice infected with recombinant BHAV expressing heterologous NSs proteins.**

IFNAR-deficient mice were mock infected or infected subcutaneously with 1 × $10^4$ PFU of recombinant BHAV variants: rIG690ΔNSs:mGL (NSs-deficient control expressing mGreenLantern), rIG690NSsV5 (IG690 NSs), ribAr2709NSsV5 (ibAr2709 NSs), or rR1819NSsV5 (R1819 NSs). **(A)** Kaplan-Meier survival curves showing the proportion of animals remaining alive over the 7-day monitoring period. **(B)** Daily clinical scoring of disease severity. Each row represents experimental groups; columns represent successive days post-infection (days 0–7). Heatmap intensity reflects cumulative clinical score (0 = no signs; 1 = mild signs; 2 = moderate signs; 3 = severe signs), with darker colours indicating more severe disease manifestations. See S2 Table for detailed scoring criteria.

| Primer Name | Primer Sequence | Purpose |
|---|---|---|
| Oligo(dT)15 primer | GACCACGCGTATCGATGTCGACTTTTTTTTTTTTTTTV | RACE - RT primer |
| PCR anchor primer | GACCACGCGTATCGATGTCGAC | RACE - Poly-A PCR primer |
| IG690 S - 5'UTR PCR | CACTCTTGTGGGAGTGATGTCG | RACE - IG690 S UTRs |
| IG690 S - 5'UTR SEQ | ATGACTTTAGCGTCATCTTTCCAG | |
| IG690 S - 3'UTR PCR | GTGCCATTCCCATACACCTTGAG | |
| IG690 S - 3'UTR SEQ | CCTTGAAATAGATCCACTAACTCC | |
| IG690 M - 5'UTR PCR | GAAGATCCCGTCCAGGATGTC | RACE - IG690 M UTRs |
| IG690 M - 5'UTR SEQ | TATCCTCTGGGATATTGACAGGC | |
| IG690 M - 3'UTR PCR | TCAGAAGGGCATCAACAATGTGAAT | |
| IG690 M - 3'UTR SEQ | GGCTTCTTAAATTGGCTAGACAC | |
| IG690 L - 5'UTR PCR | CAGGGAAATGGCGTGAGAGAG | RACE - IG690 L UTRs |
| IG690 L - 5'UTR SEQ | ACTAACACTATCTGGGACTTCATC | |
| IG690 L - 3'UTR PCR | AGAAGACTACATCAAGAGACTAGG | |
| IG690 L - 3'UTR SEQ | TGAAAACCCTGACGAGCTGTGCATACTGCCTGACAAAG | |
| IG690 S - N mRNA termination PCR | CTGACAACATCAAGGCCCTGTG | RACE - IG690 N & NSs mRNA termination signals |
| IG690 S - N mRNA termination SEQ | CTCCAGACCTTTGAGAATGCTGTG | |
| IG690 S - NSs mRNA termination PCR | TCCAAGTACTCCAAGCTTTGGGC | |
| IG690 S - NSs mRNA termination SEQ | ATTACCCTGTTAAATTGCCACG | |
| IG690 S - RT-PCR | ACACAAAGAAGCCGCGAAC | RT-PCR - IG690 S |
| IG690 S - RT-PCR | ACACAGAGAAGCCGCAGAATG | |
| IG690 M - RT-PCR | ACACAAAGACGCCAGCTATC | RT-PCR - IG690 M |
| IG690 M - RT-PCR | ACACAAAGACCGCCAGCACTG | |
| IG690 L - RT-PCR | ACACAGAGACGCCAGCATG | RT-PCR - IG690 L |
| IG690 L - RT-PCR | ACACAAAGACCGCCAGCAG | |
| ibAr2709 S - RT-PCR | ACACAAAGAAGCCGCAAAC | RT-PCR - ibAr2709 S |
| ibAr2709 S - RT-PCR | ACACAGAGAAGCCGCACC | |
| R1819, R1329 and R1336 S - RT-PCR | ACACAAAGAAGCCGCTAAC | RT-PCR - R1819, R1329 & R1336 S |
| R1819, R1329 and R1336 S - RT-PCR | ACACAGAGAAGCCGCAGAATG | |
| pTVT7 backbone - PCR F primer | GGGTCGGCATGGCATCTC | rWT Rescue transcription plasmids |
| pTVT7 backbone - PCR R primer | CTATAGTGAGTCGTATTAATTTCGCGGGATCG | |

| | | |
|---|---|---|
| IG690 S - Overhang PCR into pTVT7 F primer | TACGACTCACTATAGACACAAAGAAGCCGCGAAC | |
| IG690 S - Overhang PCR into pTVT7 R primer | ATGCCATGCCGACCCACACAGAGAAGCCGCAGAATG | |
| IG690 M - Overhang PCR into pTVT7 F primer | TACGACTCACTATAGACACAAAGACGCCAGCTATC | |
| IG690 M - Overhang PCR into pTVT7 R primer | ATGCCATGCCGACCCACACAAAGACCGCCAGCACTG | |
| IG690 L - Overhang PCR into pTVT7 F primer | TACGACTCACTATAGACACAGAGACGCCAGCATG | |
| IG690 L - Overhang PCR into pTVT7 R primer | ATGCCATGCCGACCCACACAAAGACCGCCAGCAG | |
| pCMV backbone - PCR F primer | TTTCTAGAGCGGCCGCTTC | For rWT Rescue helper plasmids |
| pCMV backbone - PCR R primer | GGTGGCTAGCCTATAGTGAG | |
| IG690 N ORF - Overhang PCR into pCMV F primer | CTCACTATAGGCTAGCCACCATGGTTGCATACACTGATATC | |
| IG690 N ORF - Overhang PCR into pCMV R primer | CGAAGCGGCCGCTCTAGAAATTACTCCAGTTTTTCCCAG | |
| IG690 L ORF - Overhang PCR into pCMV F primer | CTCACTATAGGCTAGCCACCATGGAAACTAGGATACAGG | |
| IG690 L ORF - Overhang PCR into pCMV R primer | CGAAGCGGCCGCTCTAGAAATCAACCCCAAATGTCTTC | |

**S1 Table. Oligonucleotides used in this study**

| Mild (Score 1) | Moderate (Score 2) | Advanced (Score 3) |
|---|---|---|
| Reduced weight gain | Weight loss between 10-15% | Weight loss greater than 15% |
| Slight unkempt appearance | Piloerection | Staring coat-marked piloerection |
| Subdued but responsive, animal shows normal provoked patterns of behaviour | Subdued animal shows subdued behaviour patterns | Unresponsive to extraneous activity and provocation |
| Hunched only occasionally | Hunched intermittently | Hunched persistently/markedly hunched |
| Very slight alteration in respiration | Intermittent abnormal breathing | Laboured respiration |
| Eyes slightly dull, not bright | Semi-closed eyelids | Closed eyes most of the time |
| Interacts with peers | Reduced peer interaction | Very little or no interaction with peers/dull |
| Slight gait alteration | Intermittent tremors/incoordination OR loss of one limb function | Persistent (>24 h) tremors/incoordination OR paresis OR paraplegia |
| | Transient prostration (lying stretched out)/excitement (<1 h) | Prolonged prostration (lying stretched out) / excitement (>1 h) |
| | Sporadic loose stools | Frank diarrhoea |
| | | Loss of bladder function |
| | | Self-mutilation |

**S2 Table. Scoring system for the welfare assessment of virus challenged experimental mice.**
